# Molecular exploration of host-pathogen interactions in severe *Pseudomonas aeruginosa* infection through a multi-level data integration approach

**DOI:** 10.1101/2024.10.17.618881

**Authors:** Francesco Messina, Claudia Rotondo, Luiz Ladeira, Sara Crosetti, Michele Properzi, Valentina Dimartino, Benedetta Riccitelli, Bernard Staumont, Giovanni Chillemi, Liesbet Geris, Maria Grazia Bocci, Carla Fontana

**Author notes:** Corresponding author: Carla Fontana; (CF). Francesco Messina and Claudia Rotondo contributed equally to this work. Maria Grazia Bocci and Carla Fontana contributed equally to this work.

## Abstract

Understanding host-pathogen interactions is crucial for explaining the variability in sepsis outcomes, with *Pseudomonas aeruginosa* (*PA*) remaining a significant public health concern. In this work, we explored *PA*-human host interaction mechanisms through a data integration workflow, focusing on protein-protein and metabolite-protein interactions, along with pathway modulation in affected organs during severe infections. A scoping literature review enabled us to construct a domain-based infection network encompassing pathogenesis concepts, molecular interactions, and host response signatures, providing a wide view of the relevant mechanisms involved in severe bacterial infections. Our analysis yielded a literature-based comprehensive description of PA infection mechanisms and an annotated dataset of 189 *PA*-human interactions involving 152 proteins/molecules (109 human proteins, 3 human molecules, 34 *PA* proteins, and 5 *PA* molecules). This dataset was complemented with gene expression analysis from *in vivo PA*-infected lung samples. The results indicated a notable overexpression of proinflammatory pathways and *PA*-mediated modulation of host lung responses. Our comprehensive molecular network of *PA* infection represents a valuable tool for the understanding of severe bacterial infections and offers potential applications in predicting clinical phenotypes. Through this approach combining omics data, clinical information, and pathogen characteristics, we have provided a foundation for future research in host-pathogen interactions and the mechanistic grounds to build dynamic computational models for clinical phenotype predictions.

## Introduction

Sepsis caused by multi-drug resistant pathogens remains a leading cause of mortality in intensive care units (ICU) and represents a significant public health concern (Kumar *et al*., 2024; Rello *et al*., 2017). While it is established that microbial infection outcomes depend heavily on host conditions and spatial interactions between microbes, hosts, and other microorganisms (Saarenpaa *et al*., 2023, Mu *et al*., 2023), many molecular details of these complex relationships remain unexplored. *Pseudomonas aeruginosa* (*PA*) is one of the most common pathogens for nosocomial infections, and, along with *Acinetobacter baumannii* and *Enterobacterales* resistant to carbapenem, it was listed among critical priority pathogens for WHO (WHO, 2022; 2024). The European Centre for Disease Prevention and Control (ECDC) has included *PA* in its antimicrobial resistance surveillance program (ECDC, 2022). As an opportunistic human pathogen particularly affecting Cystic Fibrosis (CF) patients, *PA*’s clinical significance stems from multiple drug resistance mechanisms, numerous virulence factors, and biofilm production capabilities, enhancing its infection and host colonization potential (Dahal *et al*., 2023). Recently, computational approaches have aided in unravelling mechanistic insights of *PA* infections. A network-assisted silico experiment allowed the identification of novel genes for virulence and antibiotic resistance, confirmed through experimental validation, showing cross-resistance against multiple drugs due to the same genes (Hwang *et al*., 2016). In another effort, a real-time deep-learning model was applied to sepsis patients aiming to estimate prognostic outcomes from early infection phases (Boussina *et al*., 2024). The model addressed baseline acuity, comorbidities, seasonal effects, and secular trends over time, unravelling the strategic significance of computational modelling to improve the clinical outcomes in sepsis patients.

Mechanistic computational modelling, omics data analysis, and clinical research have emerged as crucial tools for bridging the gap between conceptual models and clinical practice in infectious diseases (Singh *et al*., 2023, Montaldo *et al*., 2021). By structuring key pathophysiological mechanisms and identifying conceptual domains, molecular diagrams provide novel insights into biomedical knowledge (Singh *et al*., 2023, Hemedan *et al*., 2022, Ostaszewski *et al*., 2020). The value of network-based exploratory approaches was particularly evident during the COVID-19 pandemic, where rapid identification of molecular interactions between SARS-CoV-2 and human hosts became crucial (Steiner *et al*., 2024). These virus-host interactome studies revealed infection mechanisms, explained clinical manifestations (Messina *et al*., 2020, Messina *et al*., 2021, Schmidt *et al*., 2021, Lee *et al*., 2021) and enabled a timely drug repurposing process for COVID-19 treatment (Gordon *et al*., 2020, Zhou *et al*., 2023). In that context, the resulting molecular maps of disease mechanisms (e.g. a Disease Map - https://disease-maps.io/) provided biological meaning to apparently unrelated interactions, serving as a knowledge repository, representing a solid starting point for dynamic computational modelling and data analysis, capable of facilitating mechanistic understanding of complex disease processes (Ostaszewski *et al*., 2021, Niarakis *et al*., 2023).

Building upon these advances, our study presents a data integration workflow to build a molecular map of interaction between *PA* and human hosts in severe infection. Through extensive literature review, data curation, and gene expression meta-analysis, we have documented *PA* infection pathogenic mechanisms, direct protein-protein interactions (PPI), metabolite-protein interactions (MPI), and pathway activations in affected organs, organizing these findings into three conceptual domains.

## Materials and methods

### Scoping review

We conducted independent literature reviews compliant with international reference guidelines for scoping reviews (Peters *et al*., 2015). For each domain, the scoping review outcomes were processed to identify features of *PA* interactions with the host and the direct or indirect effects that they cause within the host itself.

Using a structured search string in PubMed (Supplementary Text 1), we identified 532 articles after excluding duplicates and non-English publications. We supplemented this with 27 additional articles focusing on host response to *PA* infection in both mouse models and human patients through omics data analysis. The final selection comprised 150 articles which were categorized into three interaction levels: (1) “cell interaction level”; (2) “tissue interaction level”; and (3) “organ interaction level”. Full-text articles were evaluated by the curators to define the best possible conceptual domains (Peterson, 1996). Each article was assigned a unique reference ID (SR) and documented in Table-S1.

### Conceptual domains

First, we identified the conceptual domains that organize the information obtained from the literature, providing a hierarchical model of host-pathogen interaction, following a previous experience on mapping host-pathogen interactions in the COVID-19 Disease Map project (Singh *et al*., 2023, Montaldo *et al*., 2021). Three interaction levels within the host’s system were identified: cell, tissue, and organ. For each level, we further identified conceptual domains, describing the interactions with the pathogen (Fig. 1). A comprehensive description of these interactions can be found on Supplementary Text 1, while a summary can be found below in the results section.

**Figure 1.**
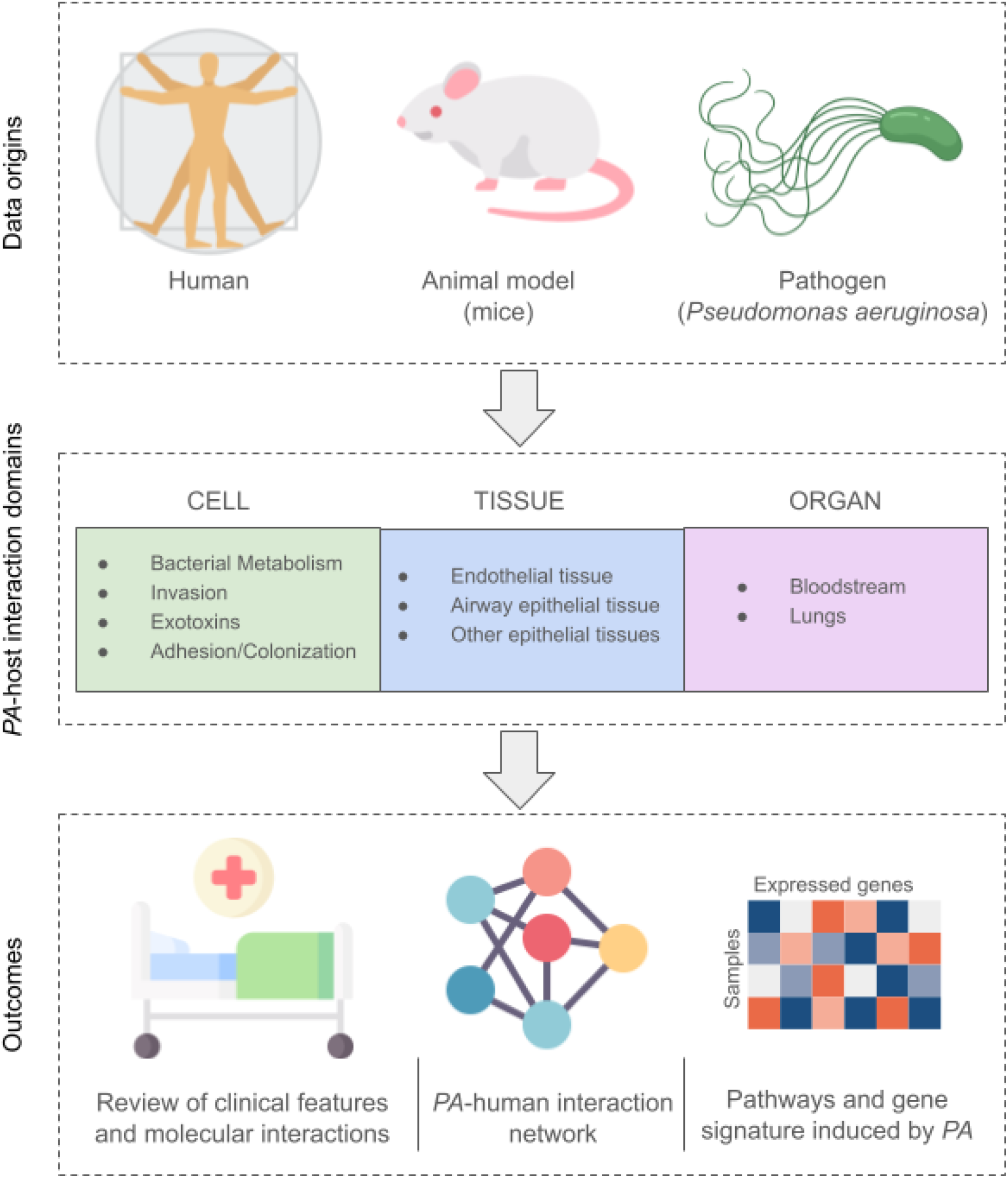
Structure of the data collection and analysis workflow. For each level, we identified further conceptual domains, describing the interactions with *PA* molecules.

### Molecular interaction dataset and human host - PA interactome

We documented PPI and MPI between *PA* and humans. All interaction details, including type, Uniprot ID, literature reference, and subdomains of the model, were compiled in the curated dataset (Table-S2). We constructed a network-based interaction model by exploring *PA*-host data gathered from the scoping review, following methodology established for SARS-CoV-2-human host interactions (Messina *et al*., 2020, Messina *et al*., 2021). Human PPI data was retrieved using R packages PSICQUIC and biomaRt (Aranda *et al*., 2011, Smedley *et al*., 2009), resulting in a comprehensive large network of 13,334 nodes and 73,584 interactions that included *PA*-human host interactions. The mechanisms of infection were estimated using the Random Walk with Restart (RWR) algorithm (Valdeolivas *et al*., 2019), using each *PA* protein as a seed and limiting the output to the 200 closest host proteins per *PA* protein. Network visualizations were generated using GEPHI\ 0.9.2 (Bastian M *et al*., 2009). Gene set enrichment analysis was performed using the R package enrichR was used to carry out gene set enrichment analysis(Kuleshov *et al*., 2016), testing against Reactome 2022, KEGG 2021 and WikiPathways 2023 human pathways databases (Milacic *et al*., 2024, Slenter *et al*., 2018, Kanehisa *et al*., 2017).

### Meta-analysis of the whole transcriptome from animal model of PA-induced sepsis

We performed a meta-analysis of gene expression in mouse lung samples comparing *PA*-infected tissues with healthy controls using data from two projects. The first dataset comprised 12 bulk RNAseq samples from *PA*-infected lung tissues (PRJNA975462; GEO: GSE233206, SRA Study SRP439193) (Yang *et al*., 2023), while the second included 6 bulk gene expression samples from acute and chronic *PA* pulmonary infection (PRJNA793679; GEO: GSE192890, SRA Study SRP353174) (Hu *et al*., 2022). SRA data was processed using Prefetch and converted to FASTQ files using the fastq-dump tool from the SRA Toolkit software v2.11.0 (Leinonen *et al*., 2011, Han *et al*., 2019). Reads were aligned to the mm10 mouse reference genome using HISAT2 (Kim *et al*., 2015). Differentially expressed (DE) genes were identified using DESeq2 v.1.42.1 in R version 3.4.3 (Love *et al*., 2014), with thresholds set at Log2FC >|1| and Benjamin-Hochberg False Discovery Rate <5% (BH-FDR). To account for batch effects between laboratories, we conducted a meta-analysis using metaRNASeq R packages, combining p-values from the two independent RNA-seq experiments using Fisher methods (Rau *et al*., 2014). The analysis focused on 21,010 genes shared between datasets, generating combined BH-adjusted p-values and average Log2FC values. Genes meeting the thresholds of Log2FC >|1| and BH FDR <5% were classified as differentially expressed genes (DEGs).

### Gene enrichment on DEGs in PA infection and healthy conditions

To deliver biological meaning from the data, we performed a gene enrichment analysis using Reactome, KEEG, and WikiPathways (Slenter *et al*., 2018, Milacic *et al*., 2024, Kanehisa *et al*., 2017). The enrichR R package was used to conduct gene set enrichment analysis, with significance assessed through Fisher exact test (p-value) and false discovery rate (q-value: adjusted p-value for FDR) (Kuleshov *et al*., 2016).

## Results

### Domain-based analysis of PA-human host interactions reveals detailed pathogenic mechanisms

At the cellular level, four key aspects characterize *PA*-host interaction: (i) bacterial adhesion/colonization (*PA*-Ad); (ii) bacterial invasion and innate immune response of the host (*PA*-In); (iii) *PA* exotoxins activity in infection (*PA*-Ex); (iv) Bacterial metabolic mechanisms (*PA*-Met). *PA* initiates infection through flagellum and type IV pili adherence, interacting with MUC1 ectodomains via NEU1 modulation (Hyun *et al*., 2016, Hyun *et al*., 2021). The bacterium employs multiple adhesion strategies, including biofilm formation, psl adhesins (Kaya *et al*., 2020, Byrd *et al*., 2010), and various receptors binding to extracellular matrix components (Paulsson *et al*., 2019, Beaufort *et al*., 2013). During invasion, *PA* modifies host cell membranes through PI3K/PIP3/Akt pathway activation and uses specialized proteins like pilY1 for binding (Engel & Eran, 2011). The bacterium’s survival in macrophages relies on mgtC and oprF (Garai *et al*., 2019) (SR22), while its exotoxins (exoS, exoT, exoU, exoY, exoA) facilitate pathogenesis through various mechanisms, including protein ribosylation, cytoskeleton modification, and membrane disruption (Henriksson *et al*., 2000, Ghatak *et al*., 2023, Jia *et al*., 2003, Krall *et al*., 2000).

At the tissue level, *PA* affects three primary domains: (i) endothelial tissue (Endothelial Tissue - EnT); (ii) lower airway and alveolar epithelial tissue in the lung, including cystic fibrosis (CF) conditions (Airway Epithelial Tissue - AET); and (iii) other epithelial tissues such as desquamated bronchial and urinary epithelia (Other Epithelial Tissues-ETs). In endothelial tissue, particularly during severe infection, APOE exhibits antibacterial activity (Puthia *et al*., 2022), while T3SS affects actin cytoskeleton dynamics (Huber *et al*., 2014). The bacterium adapts to blood survival by regulating metabolic pathways and virulence factors (Turner *et al*., 2014, Elmassry *et al*., 2019). In airway epithelial tissue, particularly relevant in CF conditions, *PA* flagella binds to asialoGM1 and MUC1, triggering inflammatory responses (Adamo *et al*., 2004, Hyun *et al*., 2021, Blohmke *et al*., 2010). CFTR plays a crucial role in *PA* uptake and inflammation (Haenisch *et al*., 2010, Schroeder *et al*., 2002). In other epithelial tissues, *PA* binds through HSPGs and N-glycans (Bucior *et al*., 2012), with quorum sensing molecules affecting barrier integrity (Vikstrom *et al*., 2009).

Finally, at the organ level, *PA* infection primarily impacts the lung and bloodstream. In lung infections, particularly in CF, *PA* causes intense inflammation with neutrophil infiltration and cytokine production (Blohmke *et al*., 2010, Vega-Carrascal *et al*., 2014). The infection involves various immune mechanisms, including TRPV4 (Scheraga *et al*., 2020), TIM3/Gal-9 signalling (Vega-Carrascal *et al*., 2014), and NET formation (Sung *et al*., 2022). Single-cell analysis has revealed significant changes in immune cell composition during infection (Hu *et al*., 2022). In bloodstream infections, *PA* affects the vascular endothelium through multiple mechanisms (Soong *et al*., 2008), with TREM-1 playing a role in septic shock (Gibot *et al*., 2019). The Hxu system contributes significantly to bloodstream infection capability (Yang *et al*., 2022a), while single-cell profiling demonstrates differential immune cell responses (Oelen *et al*., 2022). These multi-level interactions highlight the complexity of *PA* pathogenesis and its adaptive capabilities in different host environments.

**Table 1:**
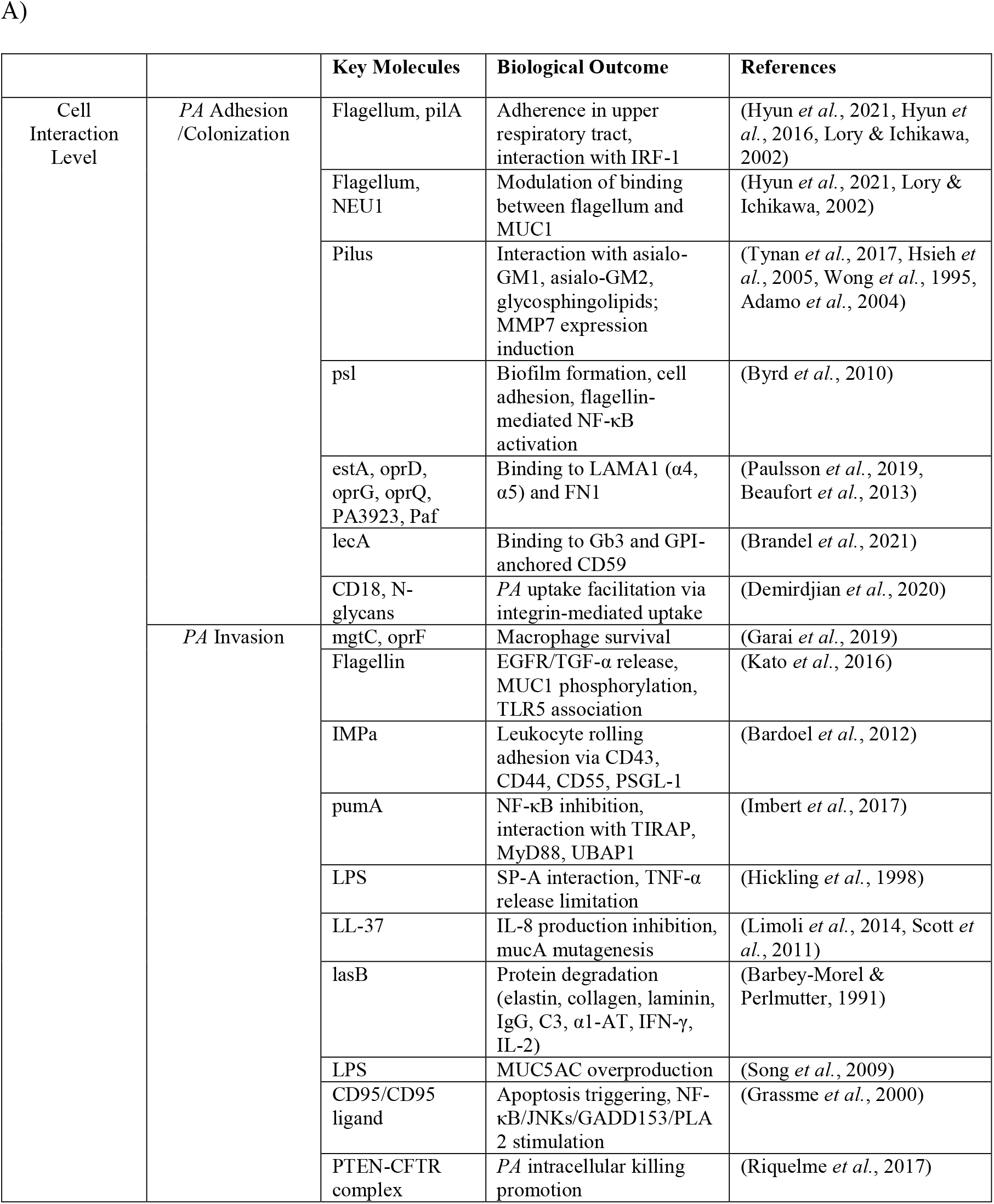
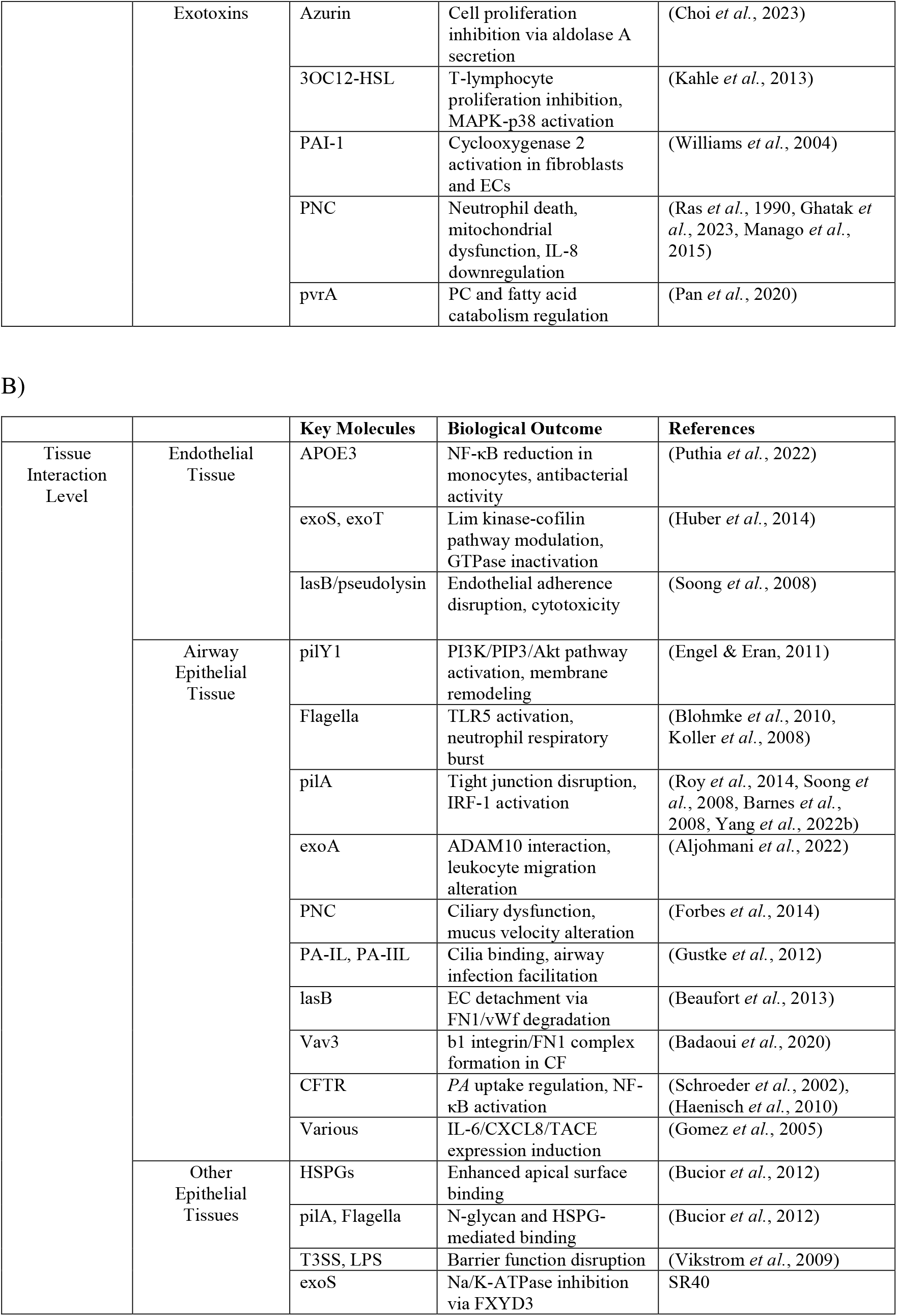
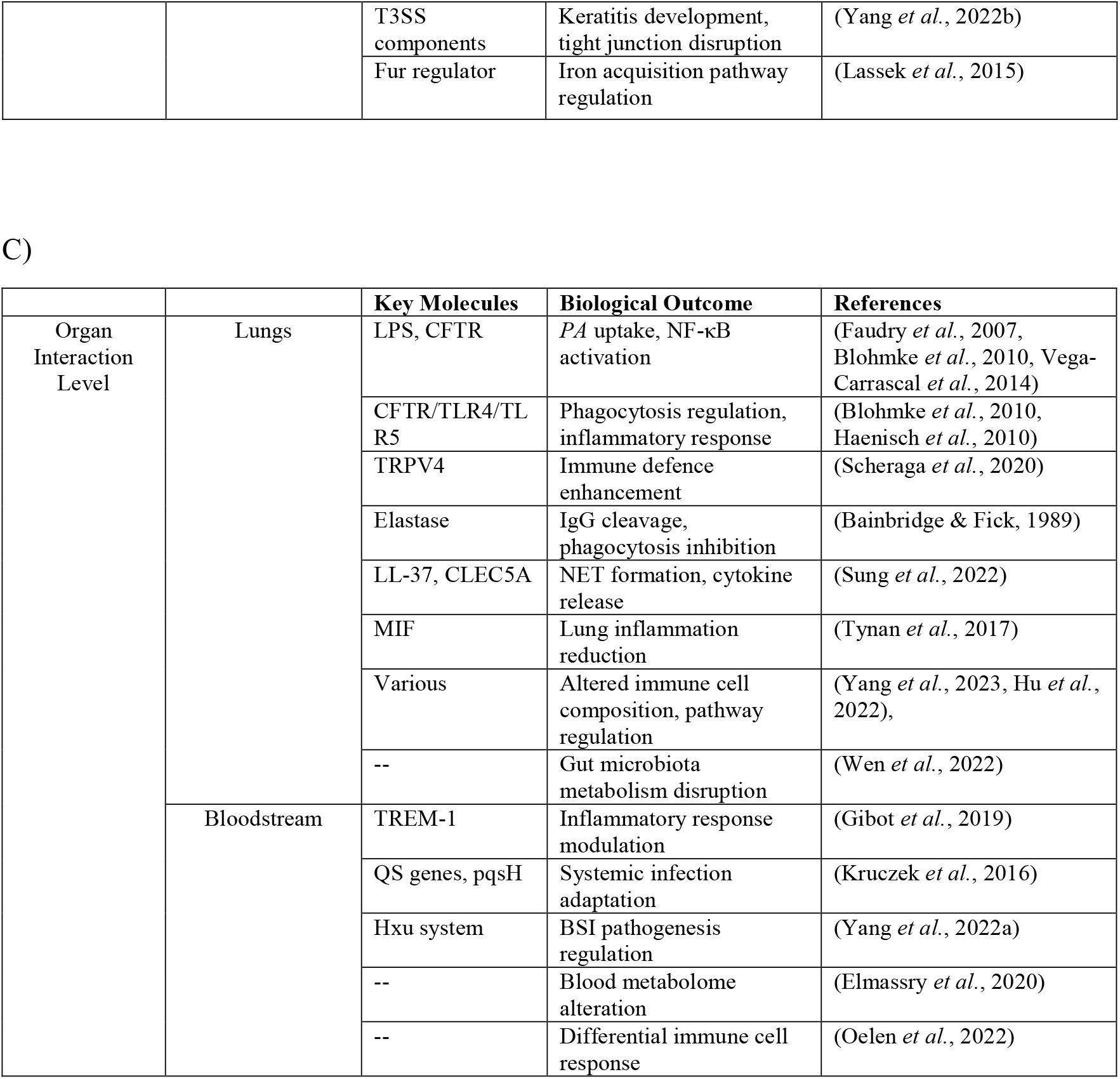
The table summarizes the main pathogenic mechanisms in *PA* infection for each domain, with comprehensive conceptual analysis provided in the Supplementary Text: A) Cell interaction level; B) Tissue interaction level; C) Organ interaction level. This structured approach enabled us to characterize specific mechanisms and experimental models of *PA* infections.

### PA-host proteins interaction network reveals key mechanisms modulated in humans by PA severe infection

We manually curated the molecular interactions between *PA* and human proteins during different infection stages. Analysis of 149 articles revealed multiple direct protein-protein interactions (PPI) and molecule-protein interactions (MPI), detailed in Table-S2 and annotated with Uniprot IDs, references, and model subdomains.

We identified 152 molecules (109 human proteins, 3 human molecules, 34 *PA* proteins, and 5 *PA* molecules), yielding 189 *PA*-human interactions and 7 human-human interactions. These interactions were categorized into four cellular domains: Adhesion process (*PA*-Ad), invasion and injury of tissue (*PA*-Inv), exotoxin production (*PA*-Ex) and bacterial metabolism (*PA*-meta).

Gene enrichment analysis revealed significant pathway associations across Reactome, WikiPathways and KEGG (Table-S3). Notable enrichments included the “Pathogenic *Escherichia coli* Infection WP2272” pathway (WikiPathway) and “Pertussis” (KEGG) with FDR<0.0001%. Reactome analysis highlighted three significant pathways (FDR<0.0001%), including Programmed Cell Death R-HSA-5357801, Toll-like Receptor Cascades R-HSA-168898, and Signalling by Interleukins R-HSA-449147. Key *PA* proteins, particularly exoS and exoT, demonstrated significant modulation of these pathways, through inhibition of interleukin proteins or degradation of occludin (OCLN), a cell death regulator (Barbieri, 2000). Notably, within the Signalling by Interleukins R-HSA-449147, we identified fibronectin (FN1), as an important player in cell adhesion, blood coagulation, and innate immunity. A full network of interactions between *PA* and human host proteins was built (Fig. 2). The network shows the overall cell response to infection, including new possible pathogenic mechanisms. The Complement Cascade Pathway (Reactome R-HSA-166658; 18/55; FDR<0.0001%) was modulated by outer membrane proteins oprH, oprQ, and the elastase lasB, contrasting cell damage by binding complement proteins like C3. These proteins also showed significant interactions with blood clotting factors (VWF, SERPINF2, PLAUR, PLAT, and PLG). GSEA analysis on the WikiPathways data highlighted the interaction with coagulation cascades (Complement and Coagulation Cascade WP558; 20/58; FDR<0.0001%), suggesting a potential involvement in thrombosis. Our network-based model also revealed a coordinated mechanism whereby exoS, exoY, and exoT trigger cell toxicity through interactions with cytoplasmic 14-3-3 proteins (e.g., YWHAB), further supporting exoS’s central role in *PA* infection.

**Figure 2.**
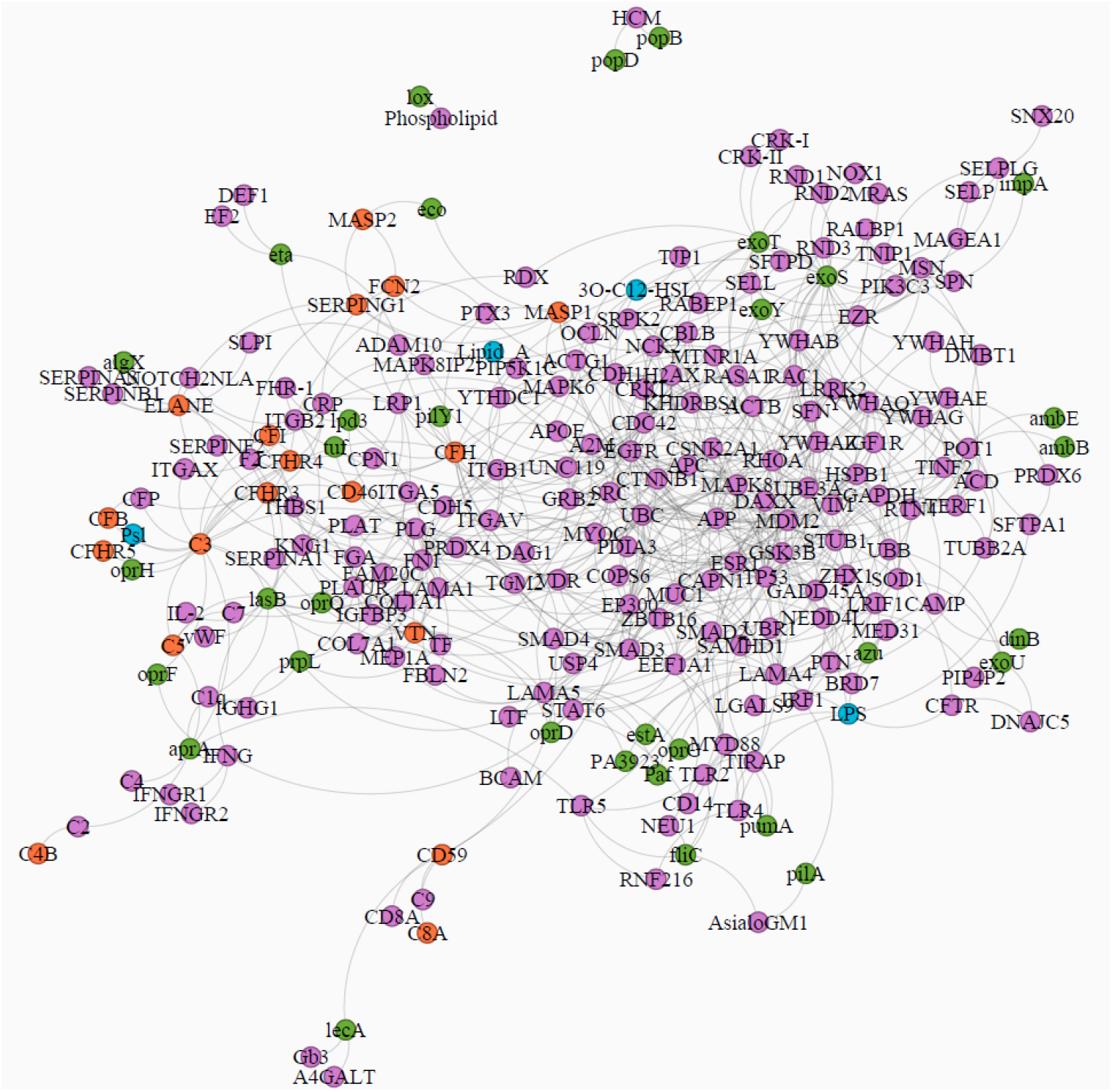
Network of *PA*-human host molecular interactions, with the top 200 nearest proteins found by the Random Walk with Restart (RWR) algorithm. Nodes have different colours to show different kinds of molecules: purple = human proteins, green = *PA* proteins, light blue = *PA* molecules, orange = human proteins belonging to the complement pathway.

### Meta-analysis of whole transcriptome of PA-infected lung tissues from mice reveals selective modulation of pro-inflammatory pathways

Our meta-analysis of gene expression of two bulk RNAseq datasets (GSE233206 and GSE192890) comparing *PA*-infected mice lung samples with healthy controls identified 1,560 upregulated and 383 downregulated genes (Log2FC > 1; FDR BH<0.05, Table S4).

Pathway analysis of upregulated genes using WikiPathways revealed significant enrichment in inflammation-related pathways, notably “Overview of Proinflammatory and Profibrotic Mediators WP5095” (39/129, FDR<0.0001%). Reactome analysis aligned with our scoping review findings, highlighting significant enrichment (FDR<0.000001%) in key pathways: Signalling by Interleukins R-HSA-449147 (103/453), Interleukin-10 Signalling R-HSA-6783783 (31/45), Cytokine Signalling in Immune System R−HSA−1280215 (145/702) (Fig. 3a/b). Proinflammatory pathways were found nested into Interleukins R-HSA-449147 Reactome’s entry (Interleukin-2 family signaling R-HSA-451927; Interleukin-3, Interleukin-5 and GM-CSF signaling R-HSA-512988; Interferon alpha/beta signaling (Homo sapiens) R-HSA-909733; Interferon gamma signaling (Homo sapiens) R-HSA-877300; ISG15 antiviral mechanism (Homo sapiens) R-HSA-1169408; PKR-mediated signaling (Homo sapiens) R-HSA-9833482; TNFR2 non-canonical NF-kB pathway (Homo sapiens) R-HSA-5668541; Signaling by CSF1 (M-CSF) in myeloid cells (Homo sapiens); R-HSA-9680350), all presenting gene targets for *PA* exoU, exoS, azu, lasB, aprA, oprF, pilA, and LPS. These results suggest that these pathways are directly involved in initiating the innate response to *PA* infection, but also highlight the potential role of *PA* molecules in modulating and limiting this response, particularly for interleukin signalling.

**Figure 3.**
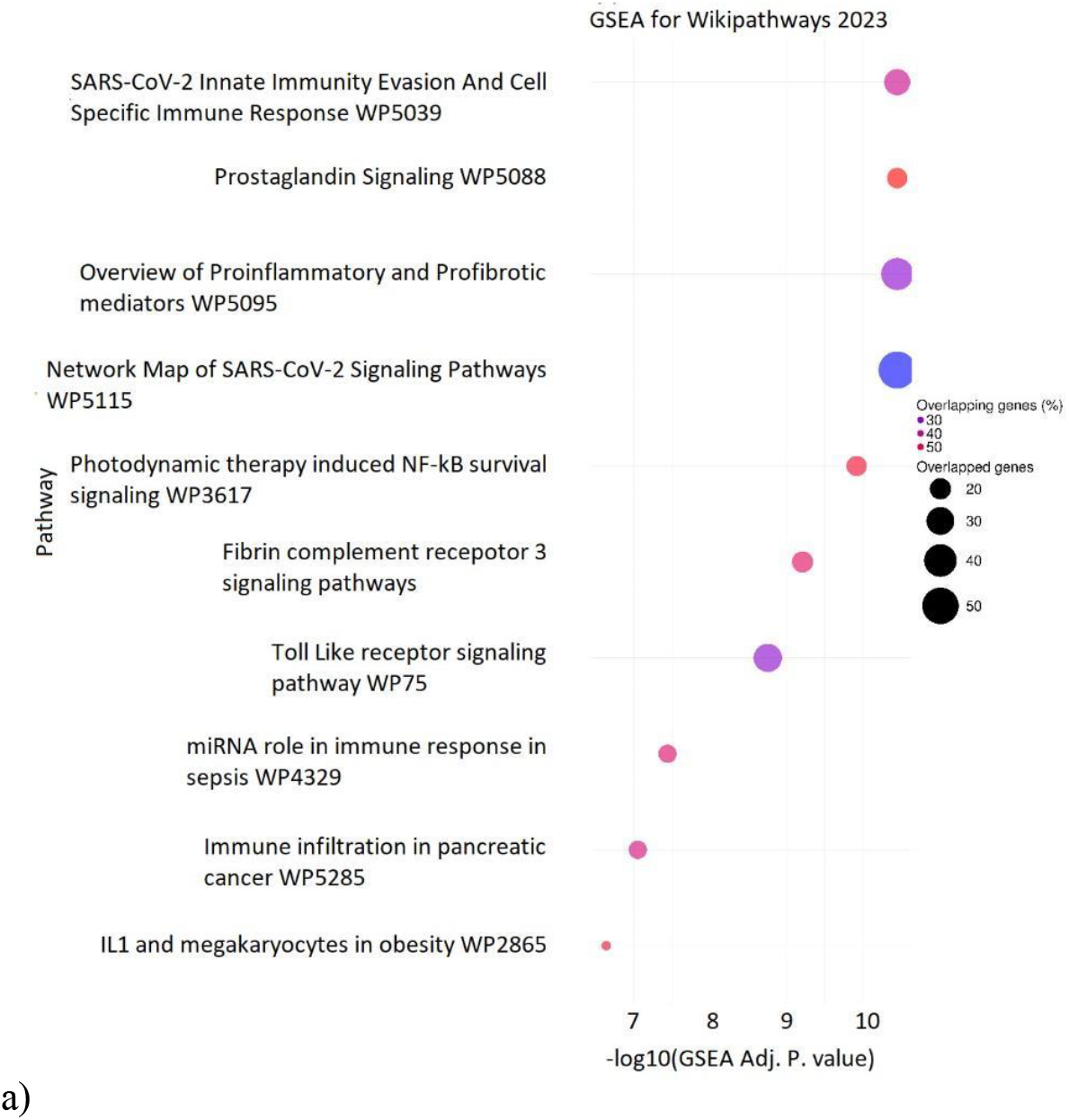
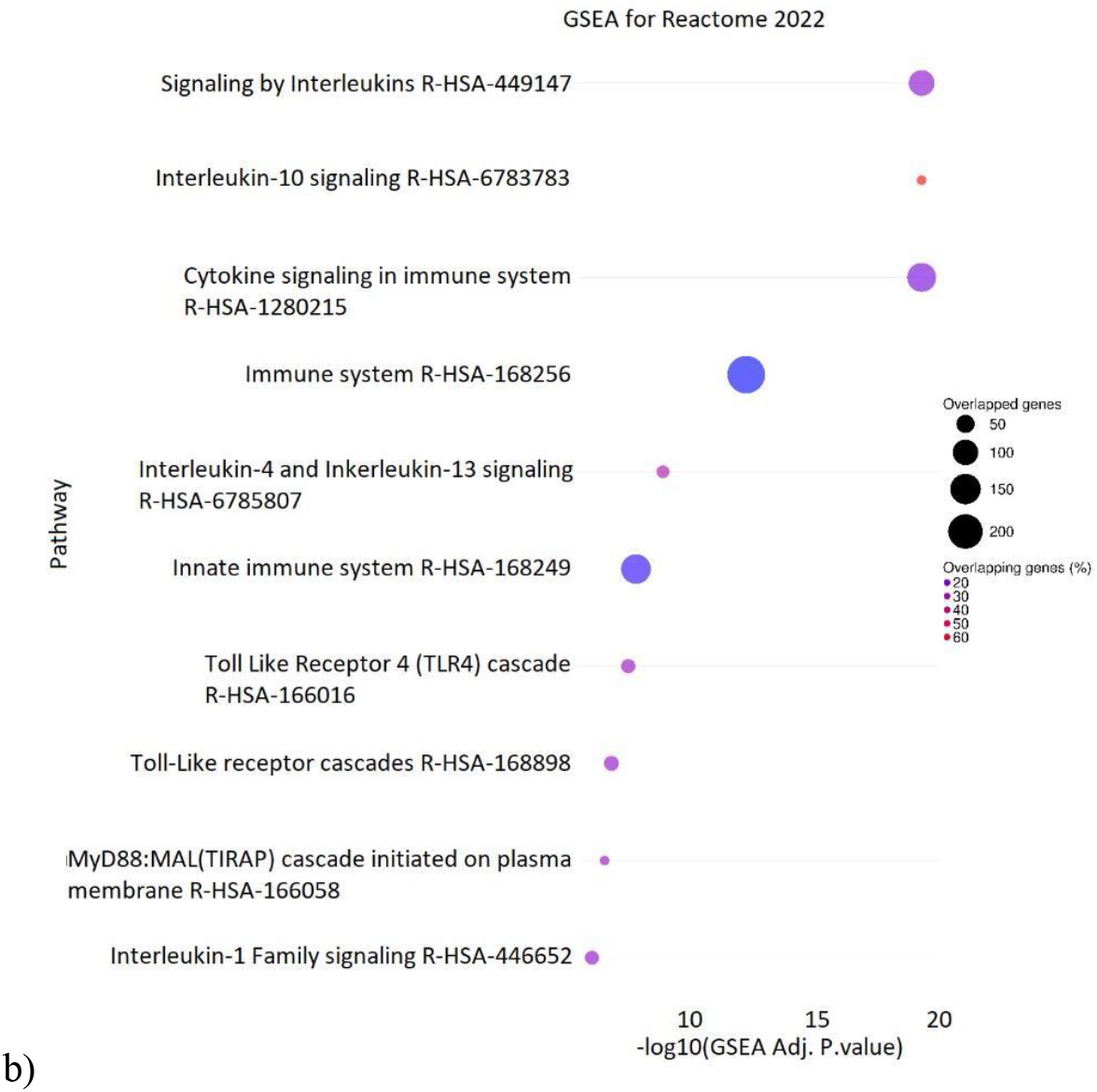
GSEA with Reactome (A) and WikiPathways (B), based on upregulated DEGs in *PA* - infected samples, obtained from meta-analysis of two infection experiments in mouse lung tissues.

## Discussion

In this work, we present our development of a comprehensive data integration model to understand *PA* infection through a detailed exploration of the literature and metanalysis of transcriptomics datasets. Our network-based explorative approach captures the complexity of severe bacterial infection by identifying specific molecular targets for each *PA* molecule, pathogenic mechanisms and host responses. *PA* serves as a complete model organism for studying severe systemic infections, given its multi-drug resistance capabilities, ability to cause acute and chronic infections in pulmonary disease patients, and its capacity to form biofilm in hypoxic conditions which makes it extremely difficult to treat [41, 42].

The computational analysis and data curation process established an annotated dataset of protein-protein interactions and molecule-protein interactions (PPI/MPI) between *PA* and human hosts, revealing several key mechanisms. Firstly, the central role of exoS during infection was confirmed, while an enhanced activity among exo family proteins, including exoY and exoT was widely highlighted (Chadha *et al*., 2022). ExoS functions by inhibiting several proteins of interleukin pathways and inducing the degradation of Occludin (OCLN), an integral membrane protein involved in cytokine-induced regulation of the tight junction permeability barrier ultimately inducing cell death (Soong *et al*., 2008). Through its ADP RT activity, exoS modulates host cell apoptosis, inducing *PA*-infected cell death by targeting various Ras proteins (Jia *et al*., 2006). The Complement Cascade Pathway undergoes modulation by *PA*’s outer membrane proteins oprH, oprQ, and elastase lasB, which trigger cytotoxic effects, and adhesion through complement protein binding, particularly C3 (Arhin & Boucher, 2010). This result mirrors the mechanism of activation of the complement system, in which C3 is the main actor against bacteria, through a link with oprF, a porin involved in ion transport (Na^+^ and Cl^−^) and anaerobic biofilm production (Sugawara *et al*., 2012, Mishra *et al*., 2015). A significant finding was the interaction between oprH, oprQ, and lasB with coagulation proteins, suggesting their activity in thrombotic processes. *PA* lasB’s cleavage of a C-terminal-peptide FYT21 derived from thrombin inhibits activation of the transcription factors NFκ-B and activator protein 1 (AP-1). *PA* demonstrates sophisticated modulation of host immune responses through multiple pathways. The bacterial proteins aprA, lasB, and exoS exhibit inhibitory effects on interleukin pathways (Chadha *et al*., 2022, Matsumoto, 2004, Phuong *et al*., 2021), indicating an adaptive modulation that enhances *PA* survival within the host. Such an effect was confirmed in *PA* infection, where *PA*-derived DnaK negatively regulates IL-1β production by cross-talk between JNK and PI3K/PDK1/FoxO1 pathways (Lee *et al*., 2020). Notably, decreased *PA* levels in CF patients correlate with reduced proinflammatory cytokines (Colombo *et al*., 2005).

Our findings provide a deep view on molecular perturbations in *PA* systemic infection and serve as a foundation for developing specific disease maps for severe *PA* infection, supporting the integration of omics data from clinical cases into predictive computational models. Future developments may incorporate text mining and AI-assisted analysis for drug target identification (Niarakis *et al*., 2023) and digital modelling of the human immune system under infection conditions (Niarakis *et al*., 2024) to better predict real patient outcomes and test potential therapeutic strategies in a personalized fashion.

Several limitations warrant consideration. While we have documented numerous significant *PA*-human interactions, our model may not encompass all possible interactions. The PPI/MPI dataset requires iterative updates to incorporate new experimental findings from both *in vitro, in vivo* and clinical studies. Furthermore, since our interaction data derives primarily from *in vitro* experiments, the described pathogenic mechanisms require validation in the context of severe systemic infections. Finally, our differential expression meta-analysis, conducted in mouse models with limited sample size, provides an overview of host gene-expression signatures in *PA* infection but requires confirmation through clinical data.

## Conclusion

Our study presents a comprehensive collection and analysis of molecular mechanisms of *P. aeruginosa* infection through a systematic data integration approach. By combining literature-based evidence, protein-protein interaction analysis, and transcriptomic data from *in vivo* studies, we have constructed a detailed dataset of *PA*-host interactions across cellular, tissue, and organ levels. Our findings highlight the complex interplay between *PA* virulence factors and host responses, particularly the role of exoS in modulating interleukin pathways and the involvement of outer membrane proteins in complement cascade regulation. The identification of 189 *PA*-human interactions involving 152 molecules provides a detailed understanding of molecular pathophysiology in invasive bacterial infections and direct molecular interactions between *PA* and host proteins, as well as involvement of molecular pathways linked to clinical phenotypes of sepsis.

The integration of differential expression analysis from mouse models further strengthens our understanding of host response patterns, particularly in proinflammatory and immune signaling pathways. As antimicrobial resistance continues to pose significant challenges in healthcare, such comprehensive molecular understanding may prove invaluable for applying precision medicine approaches to severe bacterial infections and improving patient-tailored treatments in severe systemic infections.

## Supporting information

Supplemetary text_revised

## Contributors

CF, CR and FM designed the study, developed the search strategy with feedback from LL and BS. FM, CR, VD and MP screened and selected studies. CR, VD, SC, MP and BR extracted the data and prepared the data for analysis. FM, CR, LL, BS, LG, MGB, GC and CF verified the study data. FM, LL and SC analysed the data. FM, CR, VD and MP wrote the first draft of the manuscript. FM, LL, MGB, GC and CF, writing—review and editing. MGB and CF, supervision. CF and FM, project administration; FM, LG, and CF, funding acquisition. All authors critically revised the manuscript. All authors had full access to all the data in the study and had final responsibility for the decision to submit it for publication.

## Data sharing

The datasets generated and analysed in this meta-analysis are published in the supplementary text and in supplementary tables.

## Acknowledgments

This work was supported by grants from the Italian Ministry of Health through “Ricerca Corrente” Linea 3 and ‘5 per 1000–2021’ grant of the Italian Ministry of Health (grant No. 5M-2021-23683787) (FM) and the European Commission with the HORIZON program BY-COVID (grant No. 101046203—BY-COVID). Moreover, the authors acknowledge funding from the European Union’s Horizon 2020 research and innovation programme via the European Research Council (ERC CoG INSITE 772418). Figure 1 has been designed using resources from Flaticon.com.

## References

Adamo, R., Sokol, S., Soong, G., Gomez, M.I. & Prince, A. (2004) Pseudomonas aeruginosa flagella activate airway epithelial cells through asialoGM1 and toll-like receptor 2 as well as toll-like receptor 5. Am J Respir Cell Mol Biol, 30, 627–34.

Aljohmani, A., Opitz, B., Bischoff, M. & Yildiz, D. (2022) Pseudomonas aeruginosa Triggered Exosomal Release of ADAM10 Mediates Proteolytic Cleavage in Trans. Int J Mol Sci, 23.

Aranda, B., Blankenburg, H., Kerrien, S., Brinkman, F.S., Ceol, A., Chautard, E., Dana, J.M., De Las Rivas, J., Dumousseau, M., Galeota, E., Gaulton, A., Goll, J., Hancock, R.E., Isserlin, R., Jimenez, R.C., Kerssemakers, J., Khadake, J., Lynn, D.J., Michaut, M., O’kelly, G., Ono, K., Orchard, S., Prieto, C., Razick, S., Rigina, O., Salwinski, L., Simonovic, M., Velankar, S., Winter, A., Wu, G., Bader, G.D., Cesareni, G., Donaldson, I.M., Eisenberg, D., Kleywegt, G.J., Overington, J., Ricard-Blum, S., Tyers, M., Albrecht, M. & Hermjakob, H. (2011) PSICQUIC and PSISCORE: accessing and scoring molecular interactions. Nat Methods, 8, 528–9.

Arhin, A. & Boucher, C. (2010) The outer membrane protein OprQ and adherence of Pseudomonas aeruginosa to human fibronectin. Microbiology (Reading), 156, 1415–1423.

Badaoui, M., Zoso, A., Idris, T., Bacchetta, M., Simonin, J., Lemeille, S., Wehrle-Haller, B. & Chanson, M. (2020) Vav3 Mediates Pseudomonas aeruginosa Adhesion to the Cystic Fibrosis Airway Epithelium. Cell Rep, 32, 107842.

Bainbridge, T. & Fick, R.B., Jr. (1989) Functional importance of cystic fibrosis immunoglobulin G fragments generated by Pseudomonas aeruginosa elastase. J Lab Clin Med, 114, 728–33.

Barbey-Morel, C. & Perlmutter, D.H. (1991) Effect of pseudomonas elastase on human mononuclear phagocyte alpha 1-antitrypsin expression. Pediatr Res, 29, 133–40.

Barbieri, J.T. (2000) Pseudomonas aeruginosa exoenzyme S, a bifunctional type-III secreted cytotoxin. Int J Med Microbiol, 290, 381–7.

Bardoel, B.W., Van Kessel, K.P., Van Strijp, J.A. & Milder, F.J. (2012) Inhibition of Pseudomonas aeruginosa virulence: characterization of the AprA-AprI interface and species selectivity. J Mol Biol, 415, 573–83.

Barnes, R.J., Leung, K.T., Schraft, H. & Ulanova, M. (2008) Chromosomal gfp labelling of Pseudomonas aeruginosa using a mini-Tn7 transposon: application for studies of bacteria-host interactions. Can J Microbiol, 54, 48–57.

Bastian M, Heymann S & M., J. (2009) Gephi: an open source software for exploring and manipulating networks. I.a.C.O.W.a.S. Media (ed.).

Beaufort, N., Corvazier, E., Mlanaoindrou, S., De Bentzmann, S. & Pidard, D. (2013) Disruption of the endothelial barrier by proteases from the bacterial pathogen Pseudomonas aeruginosa: implication of matrilysis and receptor cleavage. PLoS One, 8, e75708.

Blohmke, C.J., Park, J., Hirschfeld, A.F., Victor, R.E., Schneiderman, J., Stefanowicz, D., Chilvers, M.A., Durie, P.R., Corey, M., Zielenski, J., Dorfman, R., Sandford, A.J., Daley, D. & Turvey, S.E. (2010) TLR5 as an anti-inflammatory target and modifier gene in cystic fibrosis. J Immunol, 185, 7731–8.

Boussina, A., Shashikumar, S.P., Malhotra, A., Owens, R.L., El-Kareh, R., Longhurst, C.A., Quintero, K., Donahue, A., Chan, T.C., Nemati, S. & Wardi, G. (2024) Impact of a deep learning sepsis prediction model on quality of care and survival. NPJ Digit Med, 7, 14.

Brandel, A., Aigal, S., Lagies, S., Schlimpert, M., Melendez, A.V., Xu, M., Lehmann, A., Hummel, D., Fisch, D., Madl, J., Eierhoff, T., Kammerer, B. & Romer, W. (2021) The Gb3-enriched CD59/flotillin plasma membrane domain regulates host cell invasion by Pseudomonas aeruginosa. Cell Mol Life Sci, 78, 3637–3656.

Bucior, I., Pielage, J.F. & Engel, J.N. (2012) Pseudomonas aeruginosa pili and flagella mediate distinct binding and signaling events at the apical and basolateral surface of airway epithelium. PLoS Pathog, 8, e1002616.

Byrd, M.S., Pang, B., Mishra, M., Swords, W.E. & Wozniak, D.J. (2010) The Pseudomonas aeruginosa exopolysaccharide Psl facilitates surface adherence and NF-kappaB activation in A549 cells. mBio, 1.

Chadha, J., Harjai, K. & Chhibber, S. (2022) Revisiting the virulence hallmarks of Pseudomonas aeruginosa: a chronicle through the perspective of quorum sensing. Environ Microbiol, 24, 2630–2656.

Choi, J.K., Naffouje, S.A., Goto, M., Wang, J., Christov, K., Rademacher, D.J., Green, A., Stecenko, A.A., Chakrabarty, A.M., Das Gupta, T.K. & Yamada, T. (2023) Cross-talk between cancer and Pseudomonas aeruginosa mediates tumor suppression. Commun Biol, 6, 16.

Colombo, C., Costantini, D., Rocchi, A., Cariani, L., Garlaschi, M.L., Tirelli, S., Calori, G., Copreni, E. & Conese, M. (2005) Cytokine levels in sputum of cystic fibrosis patients before and after antibiotic therapy. Pediatr Pulmonol, 40, 15–21.

Dahal, S., Renz, A., Drager, A. & Yang, L. (2023) Genome-scale model of Pseudomonas aeruginosa metabolism unveils virulence and drug potentiation. Commun Biol, 6, 165.

Demirdjian, S., Hopkins, D., Cumbal, N., Lefort, C.T. & Berwin, B. (2020) Distinct Contributions of CD18 Integrins for Binding and Phagocytic Internalization of Pseudomonas aeruginosa. Infect Immun, 88.

Ecdc (2022) Antimicrobial resistance surveillance in Europe 2022. E.C.F.D.P.A. Control (ed.). Stockholm.

Elmassry, M.M., Mudaliar, N.S., Colmer-Hamood, J.A., San Francisco, M.J., Griswold, J.A., Dissanaike, S. & Hamood, A.N. (2020) New markers for sepsis caused by Pseudomonas aeruginosa during burn infection. Metabolomics, 16, 40.

Elmassry, M.M., Mudaliar, N.S., Kottapalli, K.R., Dissanaike, S., Griswold, J.A., San Francisco, M.J., Colmer-Hamood, J.A. & Hamood, A.N. (2019) Pseudomonas aeruginosa Alters Its Transcriptome Related to Carbon Metabolism and Virulence as a Possible Survival Strategy in Blood from Trauma Patients. mSystems, 4.

Engel, J. & Eran, Y. (2011) Subversion of mucosal barrier polarity by pseudomonas aeruginosa. Front Microbiol, 2, 114.

Faudry, E., Job, V., Dessen, A., Attree, I. & Forge, V. (2007) Type III secretion system translocator has a molten globule conformation both in its free and chaperone-bound forms. FEBS J, 274, 3601–3610.

Forbes, A., Davey, A.K., Perkins, A.V., Grant, G.D., Mcfarland, A.J., Mcdermott, C.M. & Anoopkumar-Dukie, S. (2014) ERK1/2 activation modulates pyocyanin-induced toxicity in A549 respiratory epithelial cells. Chem Biol Interact, 208, 58–63.

Garai, P., Berry, L., Moussouni, M., Bleves, S. & Blanc-Potard, A.B. (2019) Killing from the inside: Intracellular role of T3SS in the fate of Pseudomonas aeruginosa within macrophages revealed by mgtC and oprF mutants. PLoS Pathog, 15, e1007812.

Ghatak, S., Hemann, C., Boslett, J., Singh, K., Sharma, A., El Masry, M.S., Abouhashem, A.S., Ghosh, N., Mathew-Steiner, S.S., Roy, S., Zweier, J.L. & Sen, C.K. (2023) Bacterial Pyocyanin Inducible Keratin 6A Accelerates Closure of Epithelial Defect under Conditions of Mitochondrial Dysfunction. J Invest Dermatol, 143, 2052–2064 e5.

Gibot, S., Jolly, L., Lemarie, J., Carrasco, K., Derive, M. & Boufenzer, A. (2019) Triggering Receptor Expressed on Myeloid Cells-1 Inhibitor Targeted to Endothelium Decreases Cell Activation. Front Immunol, 10, 2314.

Gomez, M.I., Sokol, S.H., Muir, A.B., Soong, G., Bastien, J. & Prince, A.S. (2005) Bacterial induction of TNF-alpha converting enzyme expression and IL-6 receptor alpha shedding regulates airway inflammatory signaling. J Immunol, 175, 1930–6.

Gordon, D.E., Jang, G.M., Bouhaddou, M., Xu, J., Obernier, K., White, K.M., O’meara, M.J., Rezelj, V.V., Guo, J.Z., Swaney, D.L., Tummino, T.A., Huttenhain, R., Kaake, R.M., Richards, A.L., Tutuncuoglu, B., Foussard, H., Batra, J., Haas, K., Modak, M., Kim, M., Haas, P., Polacco, B.J., Braberg, H., Fabius, J.M., Eckhardt, M., Soucheray, M., Bennett, M.J., Cakir, M., Mcgregor, M.J., Li, Q., Meyer, B., Roesch, F., Vallet, T., Mac Kain, A., Miorin, L., Moreno, E., Naing, Z.Z.C., Zhou, Y., Peng, S., Shi, Y., Zhang, Z., Shen, W., Kirby, I.T., Melnyk, J.E., Chorba, J.S., Lou, K., Dai, S.A., Barrio-Hernandez, I., Memon, D., Hernandez-Armenta, C., Lyu, J., Mathy, C.J.P., Perica, T., Pilla, K.B., Ganesan, S.J., Saltzberg, D.J., Rakesh, R., Liu, X., Rosenthal, S.B., Calviello, L., Venkataramanan, S., Liboy-Lugo, J., Lin, Y., Huang, X.P., Liu, Y., Wankowicz, S.A., Bohn, M., Safari, M., Ugur, F.S., Koh, C., Savar, N.S., Tran, Q.D., Shengjuler, D., Fletcher, S.J., O’neal, M.C., Cai, Y., Chang, J.C.J., Broadhurst, D.J., Klippsten, S., Sharp, P.P., Wenzell, N.A., Kuzuoglu-Ozturk, D., Wang, H.Y., Trenker, R., Young, J.M., Cavero, D.A., Hiatt, J., Roth, T.L., Rathore, U., Subramanian, A., Noack, J., Hubert, M., Stroud, R.M., Frankel, A.D., Rosenberg, O.S., Verba, K.A., Agard, D.A., Ott, M., Emerman, M., Jura, N., et al. (2020) A SARS-CoV-2 protein interaction map reveals targets for drug repurposing. Nature, 583, 459–468.

Grassme, H., Kirschnek, S., Riethmueller, J., Riehle, A., Von Kurthy, G., Lang, F., Weller, M. & Gulbins, E. (2000) CD95/CD95 ligand interactions on epithelial cells in host defense to Pseudomonas aeruginosa. Science, 290, 527–30.

Gustke, H., Kleene, R., Loers, G., Nehmann, N., Jaehne, M., Bartels, K.M., Jaeger, K.E., Schachner, M. & Schumacher, U. (2012) Inhibition of the bacterial lectins of Pseudomonas aeruginosa with monosaccharides and peptides. Eur J Clin Microbiol Infect Dis, 31, 207–15.

Haenisch, M.D., Ciche, T.A. & Luckie, D.B. (2010) Pseudomonas or LPS exposure alters CFTR iodide efflux in 2WT2 epithelial cells with time and dose dependence. Biochem Biophys Res Commun, 394, 1087–92.

Han, Z., Hua, J., Xue, W. & Zhu, F. (2019) Integrating the Ribonucleic Acid Sequencing Data From Various Studies for Exploring the Multiple Sclerosis-Related Long Noncoding Ribonucleic Acids and Their Functions. Front Genet, 10, 1136.

Hemedan, A.A., Niarakis, A., Schneider, R. & Ostaszewski, M. (2022) Boolean modelling as a logic-based dynamic approach in systems medicine. Comput Struct Biotechnol J, 20, 3161–3172.

Henriksson, M.L., Rosqvist, R., Telepnev, M., Wolf-Watz, H. & Hallberg, B. (2000) Ras effector pathway activation by epidermal growth factor is inhibited in vivo by exoenzyme S ADP-ribosylation of Ras. Biochem J, 347 Pt 1, 217–22.

Hickling, T.P., Sim, R.B. & Malhotra, R. (1998) Induction of TNF-alpha release from human buffy coat cells by Pseudomonas aeruginosa is reduced by lung surfactant protein A. FEBS Lett, 437, 65–9.

Hsieh, J.C., Tham, D.M., Feng, W., Huang, F., Embaie, S., Liu, K., Dean, D., Hertle, R., Fitzgerald, D.J. & Mrsny, R.J. (2005) Intranasal immunization strategy to impede pilin-mediated binding of Pseudomonas aeruginosa to airway epithelial cells. Infect Immun, 73, 7705–17.

Hu, X., Wu, M., Ma, T., Zhang, Y., Zou, C., Wang, R., Ren, Y., Li, Q., Liu, H., Li, H., Wang, T., Sun, X., Yang, Y., Tang, M., Li, X., Li, J., Gao, X., Li, T. & Zhou, X. (2022) Single-cell transcriptomics reveals distinct cell response between acute and chronic pulmonary infection of Pseudomonas aeruginosa. MedComm (2020), 3, e193.

Huber, P., Bouillot, S., Elsen, S. & Attree, I. (2014) Sequential inactivation of Rho GTPases and Lim kinase by Pseudomonas aeruginosa toxins ExoS and ExoT leads to endothelial monolayer breakdown. Cell Mol Life Sci, 71, 1927–41.

Hwang, S., Kim, C.Y., Ji, S.G., Go, J., Kim, H., Yang, S., Kim, H.J., Cho, A., Yoon, S.S. & Lee, I. (2016) Network-assisted investigation of virulence and antibiotic-resistance systems in Pseudomonas aeruginosa. Sci Rep, 6, 26223.

Hyun, S.W., Imamura, A., Ishida, H., Piepenbrink, K.H., Goldblum, S.E. & Lillehoj, E.P. (2021) The sialidase NEU1 directly interacts with the juxtamembranous segment of the cytoplasmic domain of mucin-1 to inhibit downstream PI3K-Akt signaling. J Biol Chem, 297, 101337.

Hyun, S.W., Liu, A., Liu, Z., Cross, A.S., Verceles, A.C., Magesh, S., Kommagalla, Y., Kona, C., Ando, H., Luzina, I.G., Atamas, S.P., Piepenbrink, K.H., Sundberg, E.J., Guang, W., Ishida, H., Lillehoj, E.P. & Goldblum, S.E. (2016) The NEU1-selective sialidase inhibitor, C9-butyl-amide-DANA, blocks sialidase activity and NEU1-mediated bioactivities in human lung in vitro and murine lung in vivo. Glycobiology, 26, 834–49.

Imbert, P.R., Louche, A., Luizet, J.B., Grandjean, T., Bigot, S., Wood, T.E., Gagne, S., Blanco, A., Wunderley, L., Terradot, L., Woodman, P., Garvis, S., Filloux, A., Guery, B. & Salcedo, S.P. (2017) A Pseudomonas aeruginosa TIR effector mediates immune evasion by targeting UBAP1 and TLR adaptors. EMBO J, 36, 1869–1887.

Jia, J., Alaoui-El-Azher, M., Chow, M., Chambers, T.C., Baker, H. & Jin, S. (2003) c-Jun NH2-terminal kinase-mediated signaling is essential for Pseudomonas aeruginosa ExoS-induced apoptosis. Infect Immun, 71, 3361–70.

Jia, J., Wang, Y., Zhou, L. & Jin, S. (2006) Expression of Pseudomonas aeruginosa toxin ExoS effectively induces apoptosis in host cells. Infect Immun, 74, 6557–70.

Kahle, N.A., Brenner-Weiss, G., Overhage, J., Obst, U. & Hansch, G.M. (2013) Bacterial quorum sensing molecule induces chemotaxis of human neutrophils via induction of p38 and leukocyte specific protein 1 (LSP1). Immunobiology, 218, 145–51.

Kanehisa, M., Furumichi, M., Tanabe, M., Sato, Y. & Morishima, K. (2017) KEGG: new perspectives on genomes, pathways, diseases and drugs. Nucleic Acids Res, 45, D353–D361.

Kato, K., Lillehoj, E.P. & Kim, K.C. (2016) Pseudomonas aeruginosa stimulates tyrosine phosphorylation of and TLR5 association with the MUC1 cytoplasmic tail through EGFR activation. Inflamm Res, 65, 225–33.

Kaya, E., Grassi, L., Benedetti, A., Maisetta, G., Pileggi, C., Di Luca, M., Batoni, G. & Esin, S. (2020) In vitro Interaction of Pseudomonas aeruginosa Biofilms With Human Peripheral Blood Mononuclear Cells. Front Cell Infect Microbiol, 10, 187.

Kim, D., Langmead, B. & Salzberg, S.L. (2015) HISAT: a fast spliced aligner with low memory requirements. Nat Methods, 12, 357–60.

Koller, B., Kappler, M., Latzin, P., Gaggar, A., Schreiner, M., Takyar, S., Kormann, M., Kabesch, M., Roos, D., Griese, M. & Hartl, D. (2008) TLR expression on neutrophils at the pulmonary site of infection: TLR1/TLR2-mediated up-regulation of TLR5 expression in cystic fibrosis lung disease. J Immunol, 181, 2753–63.

Krall, R., Schmidt, G., Aktories, K. & Barbieri, J.T. (2000) Pseudomonas aeruginosa ExoT is a Rho GTPase-activating protein. Infect Immun, 68, 6066–8.

Kruczek, C., Kottapalli, K.R., Dissanaike, S., Dzvova, N., Griswold, J.A., Colmer-Hamood, J.A. & Hamood, A.N. (2016) Major Transcriptome Changes Accompany the Growth of Pseudomonas aeruginosa in Blood from Patients with Severe Thermal Injuries. PLoS One, 11, e0149229.

Kuleshov, M.V., Jones, M.R., Rouillard, A.D., Fernandez, N.F., Duan, Q., Wang, Z., Koplev, S., Jenkins, S.L., Jagodnik, K.M., Lachmann, A., Mcdermott, M.G., Monteiro, C.D., Gundersen, G.W. & Ma’ayan, A. (2016) Enrichr: a comprehensive gene set enrichment analysis web server 2016 update. Nucleic Acids Res, 44, W90–7.

Kumar, N.R., Balraj, T.A., Kempegowda, S.N. & Prashant, A. (2024) Multidrug-Resistant Sepsis: A Critical Healthcare Challenge. Antibiotics, 13(1), 46.

Lassek, C., Burghartz, M., Chaves-Moreno, D., Otto, A., Hentschker, C., Fuchs, S., Bernhardt, J., Jauregui, R., Neubauer, R., Becher, D., Pieper, D.H., Jahn, M., Jahn, D. & Riedel, K. (2015) A metaproteomics approach to elucidate host and pathogen protein expression during catheter-associated urinary tract infections (CAUTIs). Mol Cell Proteomics, 14, 989–1008.

Lee, J.H., Jeon, J., Bai, F., Wu, W. & Ha, U.H. (2020) Negative regulation of interleukin 1beta expression in response to DnaK from Pseudomonas aeruginosa via the PI3K/PDK1/FoxO1 pathways. Comp Immunol Microbiol Infect Dis, 73, 101543.

Lee, S., Lee, Y.S., Choi, Y., Son, A., Park, Y., Lee, K.M., Kim, J., Kim, J.S. & Kim, V.N. (2021) The SARS-CoV-2 RNA interactome. Mol Cell, 81, 2838–2850 e6.

Leinonen, R., Sugawara, H. & Shumway, M. (2011) The sequence read archive. Nucleic Acids Res, 39, D19–21.

Limoli, D.H., Rockel, A.B., Host, K.M., Jha, A., Kopp, B.T., Hollis, T. & Wozniak, D.J. (2014) Cationic antimicrobial peptides promote microbial mutagenesis and pathoadaptation in chronic infections. PLoS Pathog, 10, e1004083.

Lory, S. & Ichikawa, J.K. (2002) Pseudomonas-epithelial cell interactions dissected with DNA microarrays. Chest, 121, 36S–39S.

Love, M.I., Huber, W. & Anders, S. (2014) Moderated estimation of fold change and dispersion for RNA-seq data with DESeq2. Genome Biol, 15, 550.

Manago, A., Becker, K.A., Carpinteiro, A., Wilker, B., Soddemann, M., Seitz, A.P., Edwards, M.J., Grassme, H., Szabo, I. & Gulbins, E. (2015) Pseudomonas aeruginosa pyocyanin induces neutrophil death via mitochondrial reactive oxygen species and mitochondrial acid sphingomyelinase. Antioxid Redox Signal, 22, 1097–110.

Matsumoto, K. (2004) Role of bacterial proteases in pseudomonal and serratial keratitis. Biol Chem, 385, 1007–16.

Messina, F., Giombini, E., Agrati, C., Vairo, F., Ascoli Bartoli, T., Al Moghazi, S., Piacentini, M., Locatelli, F., Kobinger, G., Maeurer, M., Zumla, A., Capobianchi, M.R., Lauria, F.N. & Ippolito, G. (2020) COVID-19: viral-host interactome analyzed by network based-approach model to study pathogenesis of SARS-CoV-2 infection. J Transl Med, 18, 233.

Messina, F., Giombini, E., Montaldo, C., Sharma, A.A., Zoccoli, A., Sekaly, R.P., Locatelli, F., Zumla, A., Maeurer, M., Capobianchi, M.R., Lauria, F.N. & Ippolito, G. (2021) Looking for pathways related to COVID-19: confirmation of pathogenic mechanisms by SARS-CoV-2-host interactome. Cell Death Dis, 12, 788.

Milacic, M., Beavers, D., Conley, P., Gong, C., Gillespie, M., Griss, J., Haw, R., Jassal, B., Matthews, L., May, B., Petryszak, R., Ragueneau, E., Rothfels, K., Sevilla, C., Shamovsky, V., Stephan, R., Tiwari, K., Varusai, T., Weiser, J., Wright, A., Wu, G., Stein, L., Hermjakob, H. & D’eustachio, P. (2024) The Reactome Pathway Knowledgebase 2024. Nucleic Acids Res, 52, D672–D678.

Mishra, M., Ressler, A., Schlesinger, L.S. & Wozniak, D.J. (2015) Identification of OprF as a complement component C3 binding acceptor molecule on the surface of Pseudomonas aeruginosa. Infect Immun, 83, 3006–14.

Montaldo, C., Messina, F., Abbate, I., Antonioli, M., Bordoni, V., Aiello, A., Ciccosanti, F., Colavita, F., Farroni, C., Najafi Fard, S., Giombini, E., Goletti, D., Matusali, G., Rozera, G., Rueca, M., Sacchi, A., Piacentini, M., Agrati, C., Fimia, G.M., Capobianchi, M.R., Lauria, F.N. & Ippolito, G. (2021) Multi-omics approach to COVID-19: a domain-based literature review. J Transl Med, 19, 501.

Mu, A., Klare, W.P., Baines, S.L., Ignatius Pang, C.N., Guerillot, R., Harbison-Price, N., Keller, N., Wilksch, J., Nhu, N.T.K., Phan, M.D., Keller, B., Nijagal, B., Tull, D., Dayalan, S., Chua, H.H.C., Skoneczny, D., Koval, J., Hachani, A., Shah, A.D., Neha, N., Jadhav, S., Partridge, S.R., Cork, A.J., Peters, K., Bertolla, O., Brouwer, S., Hancock, S.J., Alvarez-Fraga, L., De Oliveira, D.M.P., Forde, B., Dale, A., Mujchariyakul, W., Walsh, C.J., Monk, I., Fitzgerald, A., Lum, M., Correa-Ospina, C., Roy Chowdhury, P., Parton, R.G., De Voss, J., Beckett, J., Monty, F., Mckinnon, J., Song, X., Stephen, J.R., Everest, M., Bellgard, M.I., Tinning, M., Leeming, M., Hocking, D., Jebeli, L., Wang, N., Ben Zakour, N., Yasar, S.A., Vecchiarelli, S., Russell, T., Zaw, T., Chen, T., Teng, D., Kassir, Z., Lithgow, T., Jenney, A., Cole, J.N., Nizet, V., Sorrell, T.C., Peleg, A.Y., Paterson, D.L., Beatson, S.A., Wu, J., Molloy, M.P., Syme, A.E., Goode, R.J.A., Hunter, A.A., Bowland, G., West, N.P., Wilkins, M.R., Djordjevic, S.P., Davies, M.R., Seemann, T., Howden, B.P., Pascovici, D., Tyagi, S., Schittenhelm, R.B., De Souza, D.P., Mcconville, M.J., Iredell, J.R., Cordwell, S.J., Strugnell, R.A., Stinear, T.P., Schembri, M.A. & Walker, M.J. (2023) Integrative omics identifies conserved and pathogen-specific responses of sepsis-causing bacteria. Nat Commun, 14, 1530.

Niarakis, A., Laubenbacher, R., An, G., Ilan, Y., Fisher, J., Flobak, A., Reiche, K., Rodríguez Martínez, M., Geris, L., Ladeira, L., Veschini, L., Blinov, M.L., Messina, F., Fonseca, L.L., Ferreira, S., Montagud, A., Noël, V., Marku, M., Tsirvouli, E., Torres, M., Harris, L.A., Sego, T.J., Cockrell, C., Shick, A., Balci, H., Salazar, A., Rian, K., Hemedan, H.A., Esteban-Medina, M., Staumont, B., Hernandez-Vargas, E., Martis B. S.,, Madrid-Valiente, A., Karampelesis, P., Sordo Vieira, L., Harlapur, P., Kulesza, A., Nikaein, N., Garira, W., Malik Sheriff, R.S., Thakar, J., Du T. Tran, V., Carbonell-Caballero, J., Safaei, S., Valencia, A., Zinovyev, A. & Glazier, J.A. (2024) Building an international and interdisciplinary community to develop immune digital twins for complex human pathologies. zenodo.

Niarakis, A., Ostaszewski, M., Mazein, A., Kuperstein, I., Kutmon, M., Gillespie, M.E., Funahashi, A., Acencio, M.L., Hemedan, A., Aichem, M., Klein, K., Czauderna, T., Burtscher, F., Yamada, T.G., Hiki, Y., Hiroi, N.F., Hu, F., Pham, N., Ehrhart, F., Willighagen, E.L., Valdeolivas, A., Dugourd, A., Messina, F., Esteban-Medina, M., Pena-Chilet, M., Rian, K., Soliman, S., Aghamiri, S.S., Puniya, B.L., Naldi, A., Helikar, T., Singh, V., Fernandez, M.F., Bermudez, V., Tsirvouli, E., Montagud, A., Noel, V., Ponce-De-Leon, M., Maier, D., Bauch, A., Gyori, B.M., Bachman, J.A., Luna, A., Pinero, J., Furlong, L.I., Balaur, I., Rougny, A., Jarosz, Y., Overall, R.W., Phair, R., Perfetto, L., Matthews, L., Rex, D.a.B., Orlic-Milacic, M., Gomez, L.C.M., De Meulder, B., Ravel, J.M., Jassal, B., Satagopam, V., Wu, G., Golebiewski, M., Gawron, P., Calzone, L., Beckmann, J.S., Evelo, C.T., D’eustachio, P., Schreiber, F., Saez-Rodriguez, J., Dopazo, J., Kuiper, M., Valencia, A., Wolkenhauer, O., Kitano, H., Barillot, E., Auffray, C., Balling, R. & Schneider, R. (2023) Drug-target identification in COVID-19 disease mechanisms using computational systems biology approaches. Front Immunol, 14, 1282859.

Oelen, R., De Vries, D.H., Brugge, H., Gordon, M.G., Vochteloo, M., Ye, C.J., Westra, H.J., Franke, L. & Van Der Wijst, M.G.P. (2022) Single-cell RNA-sequencing of peripheral blood mononuclear cells reveals widespread, context-specific gene expression regulation upon pathogenic exposure. Nat Commun, 13, 3267.

Ostaszewski, M., Mazein, A., Gillespie, M.E., Kuperstein, I., Niarakis, A., Hermjakob, H., Pico, A.R., Willighagen, E.L., Evelo, C.T., Hasenauer, J., Schreiber, F., Drager, A., Demir, E., Wolkenhauer, O., Furlong, L.I., Barillot, E., Dopazo, J., Orta-Resendiz, A., Messina, F., Valencia, A., Funahashi, A., Kitano, H., Auffray, C., Balling, R. & Schneider, R. (2020) COVID-19 Disease Map, building a computational repository of SARS-CoV-2 virus-host interaction mechanisms. Sci Data, 7, 136.

Ostaszewski, M., Niarakis, A., Mazein, A., Kuperstein, I., Phair, R., Orta-Resendiz, A., Singh, V., Aghamiri, S.S., Acencio, M.L., Glaab, E., Ruepp, A., Fobo, G., Montrone, C., Brauner, B., Frishman, G., Monraz Gomez, L.C., Somers, J., Hoch, M., Kumar Gupta, S., Scheel, J., Borlinghaus, H., Czauderna, T., Schreiber, F., Montagud, A., Ponce De Leon, M., Funahashi, A., Hiki, Y., Hiroi, N., Yamada, T.G., Drager, A., Renz, A., Naveez, M., Bocskei, Z., Messina, F., Bornigen, D., Fergusson, L., Conti, M., Rameil, M., Nakonecnij, V., Vanhoefer, J., Schmiester, L., Wang, M., Ackerman, E.E., Shoemaker, J.E., Zucker, J., Oxford, K., Teuton, J., Kocakaya, E., Summak, G.Y., Hanspers, K., Kutmon, M., Coort, S., Eijssen, L., Ehrhart, F., Rex, D.a.B., Slenter, D., Martens, M., Pham, N., Haw, R., Jassal, B., Matthews, L., Orlic-Milacic, M., Senff Ribeiro, A., Rothfels, K., Shamovsky, V., Stephan, R., Sevilla, C., Varusai, T., Ravel, J.M., Fraser, R., Ortseifen, V., Marchesi, S., Gawron, P., Smula, E., Heirendt, L., Satagopam, V., Wu, G., Riutta, A., Golebiewski, M., Owen, S., Goble, C., Hu, X., Overall, R.W., Maier, D., Bauch, A., Gyori, B.M., Bachman, J.A., Vega, C., Groues, V., Vazquez, M., Porras, P., Licata, L., Iannuccelli, M., Sacco, F., Nesterova, A., Yuryev, A., De Waard, A., Turei, D., Luna, A., Babur, O., et al. (2021) COVID19 Disease Map, a computational knowledge repository of virus-host interaction mechanisms. Mol Syst Biol, 17, e10387.

Pan, X., Fan, Z., Chen, L., Liu, C., Bai, F., Wei, Y., Tian, Z., Dong, Y., Shi, J., Chen, H., Jin, Y., Cheng, Z., Jin, S., Lin, J. & Wu, W. (2020) PvrA is a novel regulator that contributes to Pseudomonas aeruginosa pathogenesis by controlling bacterial utilization of long chain fatty acids. Nucleic Acids Res, 48, 5967–5985.

Paulsson, M., Su, Y.C., Ringwood, T., Udden, F. & Riesbeck, K. (2019) Pseudomonas aeruginosa uses multiple receptors for adherence to laminin during infection of the respiratory tract and skin wounds. Sci Rep, 9, 18168.

Peters, M.D., Godfrey, C.M., Khalil, H., Mcinerney, P., Parker, D. & Soares, C.B. (2015) Guidance for conducting systematic scoping reviews. Int J Evid Based Healthc, 13, 141–6.

Peterson, J.W. (1996) Bacterial Pathogenesis. In: Medical Microbiology S. Baron (ed.) Medical Microbiology. 4th ed. Galveston (TX).

Phuong, M.S., Hernandez, R.E., Wolter, D.J., Hoffman, L.R. & Sad, S. (2021) Impairment in inflammasome signaling by the chronic Pseudomonas aeruginosa isolates from cystic fibrosis patients results in an increase in inflammatory response. Cell Death Dis, 12, 241.

Puthia, M., Marzinek, J.K., Petruk, G., Erturk Bergdahl, G., Bond, P.J. & Petrlova, J. (2022) Antibacterial and Anti-Inflammatory Effects of Apolipoprotein E. Biomedicines, 10.

Ras, G.J., Anderson, R., Taylor, G.W., Savage, J.E., Van Niekerk, E., Wilson, R. & Cole, P.J. (1990) Proinflammatory interactions of pyocyanin and 1-hydroxyphenazine with human neutrophils in vitro. J Infect Dis, 162, 178–85.

Rau, A., Marot, G. & Jaffrezic, F. (2014) Differential meta-analysis of RNA-seq data from multiple studies. BMC Bioinformatics, 15, 91.

Rello, J., Valenzuela-Sanchez, F., Ruiz-Rodriguez, M. & Moyano, S. (2017) Sepsis: A Review of Advances in Management. Adv Ther, 34, 2393–2411.

Riquelme, S.A., Hopkins, B.D., Wolfe, A.L., Dimango, E., Kitur, K., Parsons, R. & Prince, A. (2017) Cystic Fibrosis Transmembrane Conductance Regulator Attaches Tumor Suppressor PTEN to the Membrane and Promotes Anti Pseudomonas aeruginosa Immunity. Immunity, 47, 1169–1181 e7.

Roy, S., Karmakar, M. & Pearlman, E. (2014) CD14 mediates Toll-like receptor 4 (TLR4) endocytosis and spleen tyrosine kinase (Syk) and interferon regulatory transcription factor 3 (IRF3) activation in epithelial cells and impairs neutrophil infiltration and Pseudomonas aeruginosa killing in vivo. J Biol Chem, 289, 1174–82.

Saarenpaa, S., Shalev, O., Ashkenazy, H., Carlos, V., Lundberg, D.S., Weigel, D. & Giacomello, S. (2023) Spatial metatranscriptomics resolves host-bacteria-fungi interactomes. Nat Biotechnol.

Scheraga, R.G., Abraham, S., Grove, L.M., Southern, B.D., Crish, J.F., Perelas, A., Mcdonald, C., Asosingh, K., Hasday, J.D. & Olman, M.A. (2020) TRPV4 Protects the Lung from Bacterial Pneumonia via MAPK Molecular Pathway Switching. J Immunol, 204, 1310–1321.

Schmidt, N., Lareau, C.A., Keshishian, H., Ganskih, S., Schneider, C., Hennig, T., Melanson, R., Werner, S., Wei, Y., Zimmer, M., Ade, J., Kirschner, L., Zielinski, S., Dolken, L., Lander, E.S., Caliskan, N., Fischer, U., Vogel, J., Carr, S.A., Bodem, J. & Munschauer, M. (2021) The SARS-CoV-2 RNA-protein interactome in infected human cells. Nat Microbiol, 6, 339–353.

Schroeder, T.H., Lee, M.M., Yacono, P.W., Cannon, C.L., Gerceker, A.A., Golan, D.E. & Pier, G.B. (2002) CFTR is a pattern recognition molecule that extracts Pseudomonas aeruginosa LPS from the outer membrane into epithelial cells and activates NF-kappa B translocation. Proc Natl Acad Sci U S A, 99, 6907–12.

Scott, A., Weldon, S., Buchanan, P.J., Schock, B., Ernst, R.K., Mcauley, D.F., Tunney, M.M., Irwin, C.R., Elborn, J.S. & Taggart, C.C. (2011) Evaluation of the ability of LL-37 to neutralise LPS in vitro and ex vivo. PLoS One, 6, e26525.

Singh, V., Naldi, A., Soliman, S. & Niarakis, A. (2023) A large-scale Boolean model of the rheumatoid arthritis fibroblast-like synoviocytes predicts drug synergies in the arthritic joint. NPJ Syst Biol Appl, 9, 33.

Slenter, D.N., Kutmon, M., Hanspers, K., Riutta, A., Windsor, J., Nunes, N., Melius, J., Cirillo, E., Coort, S.L., Digles, D., Ehrhart, F., Giesbertz, P., Kalafati, M., Martens, M., Miller, R., Nishida, K., Rieswijk, L., Waagmeester, A., Eijssen, L.M.T., Evelo, C.T., Pico, A.R. & Willighagen, E.L. (2018) WikiPathways: a multifaceted pathway database bridging metabolomics to other omics research. Nucleic Acids Res, 46, D661–D667.

Smedley, D., Haider, S., Ballester, B., Holland, R., London, D., Thorisson, G. & Kasprzyk, A. (2009) BioMart--biological queries made easy. BMC Genomics, 10, 22.

Song, K.S., Kim, H.J., Kim, K., Lee, J.G. & Yoon, J.H. (2009) Regulator of G-protein signaling 4 suppresses LPS-induced MUC5AC overproduction in the airway. Am J Respir Cell Mol Biol, 41, 40–9.

Soong, G., Parker, D., Magargee, M. & Prince, A.S. (2008) The type III toxins of Pseudomonas aeruginosa disrupt epithelial barrier function. J Bacteriol, 190, 2814–21.

Steiner, S., Kratzel, A., Barut, G.T., Lang, R.M., Aguiar Moreira, E., Thomann, L., Kelly, J.N. & Thiel, V. (2024) SARS-CoV-2 biology and host interactions. Nat Rev Microbiol, 22, 206–225.

Sugawara, E., Nagano, K. & Nikaido, H. (2012) Alternative folding pathways of the major porin OprF of Pseudomonas aeruginosa. FEBS J, 279, 910–8.

Sung, P.S., Peng, Y.C., Yang, S.P., Chiu, C.H. & Hsieh, S.L. (2022) CLEC5A is critical in Pseudomonas aeruginosa-induced NET formation and acute lung injury. JCI Insight, 7.

Turner, K.H., Everett, J., Trivedi, U., Rumbaugh, K.P. & Whiteley, M. (2014) Requirements for Pseudomonas aeruginosa acute burn and chronic surgical wound infection. PLoS Genet, 10, e1004518.

Tynan, A., Mawhinney, L., Armstrong, M.E., O’reilly, C., Kennedy, S., Caraher, E., Julicher, K., O’dwyer, D., Maher, L., Schaffer, K., Fabre, A., Mckone, E.F., Leng, L., Bucala, R., Bernhagen, J., Cooke, G. & Donnelly, S.C. (2017) Macrophage migration inhibitory factor enhances Pseudomonas aeruginosa biofilm formation, potentially contributing to cystic fibrosis pathogenesis. FASEB J, 31, 5102–5110.

Valdeolivas, A., Tichit, L., Navarro, C., Perrin, S., Odelin, G., Levy, N., Cau, P., Remy, E. & Baudot, A. (2019) Random walk with restart on multiplex and heterogeneous biological networks. Bioinformatics, 35, 497–505.

Vega-Carrascal, I., Bergin, D.A., Mcelvaney, O.J., Mccarthy, C., Banville, N., Pohl, K., Hirashima, M., Kuchroo, V.K., Reeves, E.P. & Mcelvaney, N.G. (2014) Galectin-9 signaling through TIM-3 is involved in neutrophil-mediated Gram-negative bacterial killing: an effect abrogated within the cystic fibrosis lung. J Immunol, 192, 2418–31.

Vikstrom, E., Bui, L., Konradsson, P. & Magnusson, K.E. (2009) The junctional integrity of epithelial cells is modulated by Pseudomonas aeruginosa quorum sensing molecule through phosphorylation-dependent mechanisms. Exp Cell Res, 315, 313–26.

Wen, L., Shi, L., Kong, X.L., Li, K.Y., Li, H., Jiang, D.X., Zhang, F. & Zhou, Z.G. (2022) Gut Microbiota Protected Against pseudomonas aeruginosa Pneumonia via Restoring Treg/Th17 Balance and Metabolism. Front Cell Infect Microbiol, 12, 856633.

WHO (2022) WHO publishes list of bacteria for which new antibiotics are urgently needed. World Health Organization (ed.). Basel.

WHO (2024) Pathogens prioritization: a scientific framework for epidemic and pandemic research preparedness. World Health Organization (ed.). Basel.

Williams, S.C., Patterson, E.K., Carty, N.L., Griswold, J.A., Hamood, A.N. & Rumbaugh, K.P. (2004) Pseudomonas aeruginosa autoinducer enters and functions in mammalian cells. J Bacteriol, 186, 2281–7.

Wong, W.Y., Campbell, A.P., Mcinnes, C., Sykes, B.D., Paranchych, W., Irvin, R.T. & Hodges, R.S. (1995) Structure-function analysis of the adherence-binding domain on the pilin of Pseudomonas aeruginosa strains PAK and KB7. Biochemistry, 34, 12963–72.

Yang, F., Zhou, Y., Chen, P., Cai, Z., Yue, Z., Jin, Y., Cheng, Z., Wu, W., Yang, L., Ha, U.H. & Bai, F. (2022a) High-Level Expression of Cell-Surface Signaling System Hxu Enhances Pseudomonas aeruginosa Bloodstream Infection. Infect Immun, 90, e0032922.

Yang, J.J., Tsuei, K.C. & Shen, E.P. (2022b) The role of Type III secretion system in the pathogenesis of Pseudomonas aeruginosa microbial keratitis. Tzu Chi Med J, 34, 8–14.

Yang, Y., Ma, T., Zhang, J., Tang, Y., Tang, M., Zou, C., Zhang, Y., Wu, M., Hu, X., Liu, H., Zhang, Q., Liu, Y., Li, H., Li, J.S., Liu, Z., Li, J., Li, T. & Zhou, X. (2023) An integrated multi-omics analysis of identifies distinct molecular characteristics in pulmonary infections of Pseudomonas aeruginosa. PLoS Pathog, 19, e1011570.

Zhou, Y., Liu, Y., Gupta, S., Paramo, M.I., Hou, Y., Mao, C., Luo, Y., Judd, J., Wierbowski, S., Bertolotti, M., Nerkar, M., Jehi, L., Drayman, N., Nicolaescu, V., Gula, H., Tay, S., Randall, G., Wang, P., Lis, J.T., Feschotte, C., Erzurum, S.C., Cheng, F. & Yu, H. (2023) A comprehensive SARS-CoV-2-human protein-protein interactome reveals COVID-19 pathobiology and potential host therapeutic targets. Nat Biotechnol, 41, 128–139.

