## Supplementary material for "Molecular exploration of host-pathogen interactions in severe *Pseudomonas aeruginosa* infection through a multi-level data integration approach": Supplemetary text_revised

### **Supplementary text 1**

This work is focused on studies that provide molecular interaction, and omics data, correlated to the early onset of *PA* infection, immune response, and clinical phenotypes in severe PA infection, considering *in vivo* and *in vitro* experiments and human clinical studies, focusing on the different PA features of infection and the risks of clinical progression. The literature analysis allowed us to detail cellular and molecular mechanisms specific to *PA* infection in human hosts during different phases of infection. Such data were reported in a multilevel organization, in which each level was characterized by further conceptual domains, describing the molecular interactions. Data from human and mice hosts were identified as input, while conceptual information, molecular interaction, and activated pathways in PA severe infection were considered as potential outputs. Finally, this work provided an organization of molecular knowledge within the host system, at three defined interaction levels that determine the pathophysiology of *PA*, linking with clinical phenotypes of systemic and severe infection.

During the review process, papers were evaluated in three sequential steps: (i) title relevance; (ii) abstract screening, and (iii) identification of specific pathogenic mechanisms in PA infection through full-text analysis.

Articles were collected on PubMed on 10/07/2023 using the following search string:

(("proteins"[All Fields] OR "proteinous"[All Fields] OR "proteins"[MeSH Terms] OR "proteins"[All Fields] OR "protein"[All Fields]) AND ("interact"[All Fields] OR "interactant"[All Fields] OR "interactants"[All Fields] OR "interacted"[All Fields] OR "interacting"[All Fields] OR "interaction"[All Fields] OR "interactional"[All Fields] OR "interactions"[All Fields] OR "interactive"[All Fields] OR "interactively"[All Fields] OR "interactives"[All Fields] OR "interactivities"[All Fields] OR "interactivity"[All Fields] OR "interacts"[All Fields]) AND ("pseudomonas aeruginosa"[MeSH Terms] OR ("pseudomonas"[All Fields] AND "aeruginosa"[All Fields]) OR "pseudomonas aeruginosa"[All Fields]) AND ("hum cell"[Journal] OR ("human"[All Fields] AND "cell"[All Fields]) OR "human cell"[All Fields]))

Each interaction level comprised multiple conceptual domains, which served as conceptual repositories summarizing biological pathways, molecular interactions, and pathogenic mechanisms. Through the workflow, we generated three key outputs: a) a review of clinical features of severe *PA* infection; b) an up-to-date dataset on molecular interactions between PA and human proteins/metabolites in complex biological processes; and c) a comprehensive list of well-defined biological pathways from open-access databases for each interaction level. These outputs will contribute to developing a disease map and dynamic models to understand molecular dynamics in PA-induced sepsis, potentially shedding light into unclear pathological mechanisms and promising therapeutic strategies.

### A) *Cellular interaction level: cellular and molecular mechanisms in PA-host interactions during different phases of infection*

1. **PA Adhesion/Colonization (PA-Ad)**

The first step of PA invasion and spreading into host is adhesion and colonization of epithelial cells (ECs) (PA-Ad). PA flagellum, composed of flagellins, and the type IV pili (pilA), formed by pilin proteins (SR79), play an important role for PA adherence in the upper respiratory tract (SR10; SR73). In A549 cells exposed to PA, there is evidence of interaction between pilA and IRF-1 (SR83). Flagellum enables the binding of ectodomains of MUC1, and such binding is modulated by the enzyme NEU1 (SR10; SR73). Flagellin interacts with mouse corneal ECs, and in human CEPs it could behave similarly (SR41). The candidate receptors for the pilus-specific interactions are glycosphingolipids, such as asialo-GM1 and asialo-GM2, sialic acid containing glycosphingolipids, lactosylceramide, and glycoproteins (SR39; SR47; SR79). PA biofilm can also promote adhesion by hindering immune activity (SR1). The psl, a mannose-rich polysaccharide adhesin involved in PA-Ad and biofilm formation, is required for adherence to human cells and facilitates flagellin-mediated NF-κB activation (SR31). PA receptors estA, oprD, oprG, oprQ, PA3923, and paf have been shown to bind components of extracellular matrix, such as laminin alpha subunit 4 and 5, as well as fibronectin (SR33, SR56). In human cells, the glycosphingolipid receptor globotriaosylceramide (Gb3) is bound by PA virulence factor lecA, along with the GPI-anchored protein CD59 (SR14). PA binds annexin II (ANXA2) on human ECs through a further bond to phospholipids, such as phosphatidylserine or phosphatidylinositol (SR27). From the point of view of immunity, CD18 expression, which has been seen both in human monocytes and murine neutrophils (SR2), is required for uptake of PA but not the initial attachment of the bacteria to the host cells. N-glycans serve as cell surface ligands to facilitate integrin-mediated uptake of PA (SR2).

2. **Invasion (PA-In)**

During the invasive process, PA’s flagellin seems to induce the release of EGFR (epidermal growth factor receptor) ligand and TGF-α, along with tyrosine phosphorylation of the MUC1 cytoplasmic tail, allowing association with Toll like receptor 5 (TLR5) (SR30). To enhance the entry process into the mucosal barrier, inducing an advantageous microenvironment for invasion, PA can modify the apical cell membrane into the basolateral membrane through activation of the PI3K/PIP3/Akt pathway, which promotes remodeling of the apical membrane. Binding to the intact pseudostratified epithelium required the surface pilin associated protein pilY1 (SR21). PA can survive inside macrophages thanks to the bacterial factors mgtC and the outer membrane porin oprF (SR22). PA immunomodulating metalloprotease (IMPa) promotes the collective rolling adhesion of leucocytes, thanks to its proteolytic activity towards CD43, CD44, CD55, and PSGL-1, four glycosylated human leucocyte surface receptors, and its interaction with selectins (SR25). Quorum-sensing (QS) signal molecules, in particular 2-(2-hydroxyphenyl)-thiazole-4-carbaldehyde (IQS), modulate virulence gene expression in PA (SR19). TLR adaptors TIRAP and MyD88, as well as UBAP1, interact with the virulence factor pumA. At the same time, the virulence factor pumA inhibits NF-κB. This impedes cytokine production and TLR receptor signaling, showcasing a novel strategy for evading the innate immune response. (SR34).

LPS participates in the invasiveness of PA: human surfactant protein A (SP-A) interacts with PA LPS, preventing TNF-α release (SR43). LL-37, an antimicrobial peptide (AMPs), inhibits PA LPS-induced IL-8 production. The low concentration of LL-37 potently inhibited the formation of PA biofilm, especially in CF lung (SR53). Finally, LL-37 induces mutagenesis of mucA, leading to mucoid conversion (SR52). PA’s elastase degrades on human PBMC, connective tissue matrix proteins, including elastin, collagen, laminin, serum proteins including IgG, C3, and α1-AT, as well as the lymphokines interferon-gamma and IL-2. Further, elastase induces structural rearrangement of endogenous α1-AT during proteolytic inactivation (SR85). Elastase interacts with alginate's uronic acid units, which weakly inhibit its hydrolysis of peptides and elastin. Alginate affects lung protease inhibitors, reducing the elastase-alpha 1-proteinase inhibitor association rate and increasing the elastase- secretory leukoprotease inhibitor (SLPI) association rate (SR17).

In a mouse model, LPS from PA induces MUC5AC overproduction (SR68). The CD95/CD95 ligand system triggers apoptosis, protecting the host from PA infection. In fact, apoptosis of infected cells results in the targeting of PA into apoptotic bodies, which are rapidly internalized by other cells, and PA–mediated activation of CD95 might stimulate NF-kB, c-Jun NH2-terminal kinase, GADD153, and phospholipase PLA2, or may interfere with the functions of growth factor receptors, resulting in the secretion of defensins and/or cytokines (SR69). In a mouse model, immune PTEN-PI3K regulation depends on both the presence of CF transmembrane conductance regulator (CFTR) on the cell surface and its interaction with the tumor suppressor PTEN, while in human immune cells, PTEN, complexed with CFTR, promotes intracellular PA killing (SR49). This interaction results in reduced PA-mediated inflammation and diminished intracellular bacterial survival in mice. MIF, a key proinflammatory cytokine implicated in the pathogenesis of inflammatory diseases, seems to reduce pulmonary inflammation after PA infection in mice (SR39).

3. **Exotoxins (PA-Ex)**

PA exotoxins, exoS, exoT, exoU, exoY, and exoA, facilitate the pathogenesis of the bacterium. exoS can ribosylate many proteins, such as Ras, inducing the phosphorylation of the MAP kinase, ERK1/2 as well as PKB/Akt kinase. Further, exoS blocks the activation of Ras, as well as Rho-family GTPases, ERM proteins, and vimentin, upon stimulation with EGF, inhibiting the exchange of GDP from GTP affecting Ras activation, locking downstream signaling pathways (SR82; SR87). The exoS inhibits antiapoptotic pathways controlled by ERK1/2 and possibly p38, inducing the activation of proapoptotic signals. Additionally, PA infection upregulates CD95L/CD95 on lung ECs, leading to apoptosis, and exoS modulates host cell signaling to dictate apoptotic outcomes. In addition, the involvement of CD95L/CD95 suggests a link between exoS and JNK activation, and the involvement of CD95L/CD95 could provide a clue for linking exoS to JNK activation (SR54). Another exoS antiapoptotic pathway, seen in cell line models, involves Fas-associated death domain (FADD): exoS could cause FADD aggregation/activation with the host’s cell, modulating the PA-induced apoptosis process (SR46). Like exoS, exoT contains GAP activity toward Rho, Rac, and CDC42, causing reversible destruction of the actin cytoskeleton, leading to cell rounding, cell detachment, inhibition of cell migration, phagocytosis, and cytokinesis (SR116). The ART domains of exoT and exoS induce cellular toxicity after association with 14-3-3 proteins (SR8). exoT targets the adaptor molecules CRK and CRKL, and its activity causes rearrangement of the actin cytoskeleton. In mouse models, exoT, complexed with its substrates Crk and Cbl-b, is less stable to proteasomal degradation (SR8). Crk appears to be required for the interaction of exoT and Cbl-b that leads to rapid degradation of exoT (SR55). Further, exoT is ubiquitinated and degraded by proteasome, in a Cbl-b–dependent manner, upon E3 ubiquitin ligase activity (SR55). exoU acts on plasma membrane lipids and requires a host chaperone both to reach the plasma membrane, and to start the toxic activity in human cells. The chaperone DNAJC5 promotes exoU toxic activity (SR7). The PLA2 activity of exoU induces the release of arachidonic acid from cell membranes in culture and a subsequent increase in prostaglandin production. exoU upregulates the transcription factor c-Fos, allows the phosphorylation of c-Jun, increasing IL-8, and modulates the expression of genes for inflammation and cell death (SR23; SR78). The exoA catalyzes the ADP-ribosylation of the eukaryotic eEF-2, affecting the protein synthesis of the host cells (SR9). HNP-1, a human defensin that binds to the cell wall precursor Lipid II, could neutralize bacterial toxins, including exoA (SR16). The PA translocators popD and popB, two toxic effector proteins associated with the secretion system T3SS, interact with lipid bilayers and form pores (SR60).

4. **PA Metabolisms (PA-Met)**

The following is a description of some mechanisms triggered by PA that impact the target cell from a metabolic point of view. Azurin exerts potent cytotoxicity towards host cells, inhibiting cell proliferation by increasing the secretion of aldolase A (SR127). PA-secreted azurin can inhibit cell proliferation and tumor growth stimulating cancer cells to upregulate aldolase A secretion. 3OC12-HSL inhibits the proliferation of T-lymphocytes, modulates the cytokine synthesis in *in vitro* models, and induces the activation of MAPK-p38, which results in the activation of MAPK2 and its target LSP1, which interacts with F-actin (SR64). Focusing on QS, the autoinducer PAI-1 activates genes such as for cyclooxygenase 2 in human fibroblasts and ECs (SR48). Pyocyanin (PNC) induces rapid death of wild-type mice neutrophils, which is due to the activation of Asm, an enzyme that stabilizes pH levels in the mitochondrion, and the generation of ROS in mitochondria. In human keratinocytes, PNC alters mitochondrial function, causing a decline in NADPH and ATP levels (SR87). In mice, PNC induces neutrophils to enhance the release of myeloperoxidase and lysozyme (SR65). Moreover, PNC could induce the formation of ceramide, causing cell death, and downregulate IL-8 secretion (SR24). Of interest, pvrA coordinately regulates the genes involved in phosphatidylcholine (PC) and long chain fatty acid catabolism, the principal components of human lung surfactant. PC is also one of the major nutrient sources for PA during lung infection (SR103).

### B) *Tissue interaction level: biological response in specific human tissues and in animal models during PA infection*

1. **Airway epithelial tissue (AET)**

The identification of the host’s receptors suitable for PA binding and entry is the main motivation in studying the first steps of PA-host interaction in specific tissues. PA flagella’s protein complex, once bound to gangliotetraosylceramide containing the GalNAcb1-4Gal sequence (asialoGM1) located on the surface of airway cells, induces the expression of Matrilysin (MMP7), a metalloproteinase involved in host defense and epithelial repair (SR70). Moreover, PA flagella’s protein complex can also bind the ectodomain of MUC1: this binding, modulated by NEU1, allows the MUC1 desialylation, enhances PA flagella adhesiveness, and alters mucus viscosity in the host. Interestingly, NEU1 expression is increased in the lungs of patients with idiopathic pulmonary fibrosis (IPF) (SR10; SR74; SR35; SR51). PA flagella induce activation of TLR5 in macrophages that is required to trigger the proinflammatory cytokine that has been seen in CF AET cells following exposure to PA. Furthermore, IL-6 and IL-8, with the concomitant CXCR1 activation, could cause a respiratory burst in neutrophils, that migrated in AET (SR74; SR66). In injuries, glycolipid molecules work as receptors for pilA and provide the initial host-pathogen attachment, maybe causing an alteration of epithelial tight and adherent junctions in the AET cells, allowing bacterial invasion (SR37; SR57). Further, the pilA activates IRF-1, the gene encoding Interferon regulatory factor 1, in AET cells inducing a fast cytotoxic effect (SR63; SR84). Extracellular matrix (ECM) components and integrin receptors could contribute to the triggering of PA infection in AET. It has been shown in the A549 cell line model, in which fibronectin (FN1) and α5β1 integrins seem to be involved in PA adherence and invasion (SR71). LPS, the main virulence factor and inducer of inflammation, during PA infection, promotes MUC5AC overproduction and affects the release of ATP extracellularly, propagating inflammatory signaling to neighboring cells (SR68). The exoA interacts with epithelial ADAM10: this binding stimulates transepithelial leukocyte migration, alters protein permeability and epithelial regeneration and integrity (SR15). Another essential component in the pathology of pseudomonal lung disease is PNC: which is crucial in pseudomonal lung disease, causing reversible ciliary dysfunction in human airway cells, and decreasing tracheal mucus velocity *in vivo* (SR86). PA produces two carbohydrate-binding lectins, named PA-IL and PA-IIL, regulated by QS, which bind to the glycocalyx, including that of the airway cilia and, thus, facilitate airway infection, contributing to adhesion by binding to cilia (SR20). Elastase LasB induces EC detachment and death (anoikis) by proteolyzing FN1, degrading vWf, and inter-endothelial junctional proteins (SR56). PA stimulates the production of IL-6 in 16HBE airway ECs, induces the expression of CXCL8, and TACE transcription. TACE has anti-inflammatory effects on airway cells by regulating responsiveness to TNF-α, which is enhanced in the airways in response to bacterial infection (SR77). Deficiency of Tet2 in bronchial ECs reduced mRNA expression of tight junction (TJ) protein 1, Cxcl1, Cxcl2, Ccl20, and occludin in bronchial brushes, which are associated with perturbed barrier function at the early stage of PA pneumonia. Respiratory ECs are required for adequate host defense during PA infection and are a major cellular source for Cxcl1 and Cxcl2, which are important for recruiting neutrophils to the bronchoalveolar space (SR32). In CF condition, Vav3 leads the ectopic b1 integrin/FN1 complex, modulating the predisposition of AET to PA adhesion through cytoskeleton remodeling and its scaffold function (SR29). An important element to shed light on PA infection’s recurrence in CF is the role of the receptor CTFR, located at the apical membrane of AET cells. CTFR results to be employed in the PA uptake mechanism and in PA internalization. The CFTR-PA interaction also leads to rapid NF-kB nuclear translocation, promoting inflammation (SR59). In macrophages of healthy subjects, CFTR works with TLR4 as a receptor for phagocytosis of PA. The genetic loss of CTFR in CF patients allows AET disruption (SR58).

2. **Endothelial tissue (EnT)**

Finally, in EnT, APOE, mostly the APOE3 isoform, exhibits anti-bacterial activity against PA through aggregation ability and antimicrobial activity. Further, APOE reduced NF-κB activation in human monocytes *in vitro* and in mice *in vivo* (SR5). In an experimental murine model, T3SS affects actin cytoskeleton dynamics in endothelial cells. The exoS and exoT promote actin filament severing via the Lim kinase-cofilin pathway and inactivate RHO, RAC, and CDC42 GTPases, inhibiting membrane ruffling and filopodia, and causing stress fiber collapse and focal adhesion disruption. Cofilin activation was also observed in a mouse model of PA-induced acute pneumonia (SR62). Performing RNA sequencing (RNA-seq) and transposon-junction sequencing (Tn-seq) in two non-diabetic murine wound infection models, one acute and one non-lethal chronic infection model, it was found that PA down-regulates lipopolysaccharide O antigen biosynthesis during infection, while up-regulates genes responsible for the biosynthesis of the siderophores pyochelin and pyoverdine, and several genes with putative rules in chemotaxis (SR91). A similar study, performed in different types of trauma patients with injuries other than burns, highlights that the upregulated genes are linked to metabolic pathways, like ABC transporters, biosynthesis of secondary metabolites, two-component systems, and glycerophospholipid metabolism. These genes are differentially expressed in response to growth of PA strain UCBPP-PA14 in trauma patients' blood. Of interest, PA adapts itself not only to blood survival, but also to the presence of therapeutic molecules in the bloodstream of these patients, such as mannitol and zinc, showing the ability of PA to adjust its virulence to the situation, increasing its virulence to host macrophages while conserving its response in an environment lacking competition (SR95).

Finally, PA alters the endothelium, producing extracellular proteinases that are also virulence factors. In human PA infections, proteinases, particularly metalloproteinases and serine-proteinases, are notable virulence factors. lasB/pseudolysin, a member of the thermolysin/M4 superfamily of metallopeptidases, is emphasized among these. This superfamily includes proteinases secreted by various common pathogenic bacteria that affect the bloodstream (SR56). This protein can severely affect the adherence and thus the survival of human ECs in culture, and the high lasB producer strain PAO1 appears to have a cytotoxic effect (SR57).

3. **Other epithelial tissues (ETs)**

PA strongly adheres to many epithelial tissues, not only bronchial ECs. In incompletely polarized cells, a model for tissue injury, chains of heparan sulfate proteoglycans (HSPGs) are upregulated at the apical (AP) surface, leading to enhanced PA binding and subsequent tissue damage (SR44). pilA mediates binding and entry at the AP surface through N-glycans, while flagella’s protein complex is required to mediate maximal binding and entry through HSPGs at the basolateral (BL) surface of polarized epithelium, although they can also mediate bacterial binding to N-glycans at the AP surface of polarized epithelium. Further, the flagellar cap protein of the only strain PAO1 binds to Lewis X oligosaccharides found at the periphery of human respiratory mucins (SR44). In the intestinal epithelium, lasI and rhlI, two proteins of QS, produce the autoinducers homoserine lactone molecules (3O-C12-HSL) and C4-HSL, respectively. 3O-C12-HSL can disrupt barrier integrity in human epithelial Caco-2 cell line through the reduction of the expression, distribution, and phosphorylation of zonular occludin 1 (ZO-1), β-catenin, E-cadherin, and occludin and the reorganization of F-actin. T3SS and LPS can affect epithelial barrier function (SR61). Focusing on T3SS, exoS binds FXYD3, a mammalian factor expressed largely in the colon and stomach that regulates Na/K-ATPase. This bind leads to the inhibition of Na/K-ATPase causing an increase in TJs permeability against ionic and nonionic solutes (SR40). Moreover, T3SS, along with exoS, exoU, exoT, and exoY, have been associated with PA keratitis, enable the invasiveness of PA through the breakdown of TJs, and affect pseudomonal adherence to contact lenses (SR84). A specific PA proteomic profile was detected exclusively in real infections, where iron acquisition pathways would be upregulated. The expression of these proteins is dependent on the PA iron-binding Fur regulator. PA proteins found exclusively in the urinary tract are secreted virulence factors involved in host-pathogen interactions (SR50). Of interest, the global urinary metabolic profile, obtained through liquid chromatography based on mass spectrometry, approaches for untargeted metabolomics, accurately discriminates ventilator-associated pneumonia (VAP) due to *PA* or other aetiologies (SR101). Shotgun quantitative proteomics analysis of proteins expressed in PA clinical isolates derived from different infection sites shows a site-specific proteome. PA isolates from the urinary tract express about three times more proteins that participate in iron and amino acid transport compared to CF-lung isolates. Interestingly, all isolates from burned wound infections exhibit elevated levels of the outer membrane protein oprH, which is involved in PA antibiotic resistance, and of the response regulator phoP, which expression is induced by low extracellular magnesium concentrations, compared to lung/urinary tract isolates. Taken together, PA has evolved a metabolic adaptation to different host environments (SR102).

### C) *Organ interaction level: response mechanism in organ injuries and systemic responses in PA infection*

1. **Lung**

PA infection causes chronic lung infections in individuals with the genetic disease CF, which is an autosomal recessive disorder resulting from mutations in the CTFR gene encoding a chloride channel. Inflammation in the CF lung is characterized by an intense neutrophilic infiltrate, high levels of proinflammatory cytokines, chemokines and evidence of activation of NF-kB and other proinflammatory signaling cascades (SR74), along with chronic bacterial infection and inflammation in the lower airways (SR76). PA LPS is specifically recognized and bound by CFTR: this binding allows internalization of the pathogen and subsequent activation of the transcription factor NF-kappa B (SR60). CFTR works as a receptor for phagocytosis of PA, together with TLR4 (SR58), and the activation of TLR5 by flagellin is required to trigger the exaggerated proinflammatory cytokine production seen in CF airway cells following exposure to PA (SR74). In CF, along with PA colonization, Vav3, a protein regulating the cytoskeleton and signal transduction, is overexpressed in primary human airway ECs (SR29). Furthermore, TRPV4 gene enhances host innate immune defenses and protects the lung from injury after PA pneumonia by enhancing macrophage bacterial clearance. It downregulates proinflammatory cytokine secretion due to modulating LPS-TLR4 signal pathway via MAPK activation switching by DUSP1 (SR11). Chronic inflammation in CF and bacterial clearance depend on the binding of gal 9 to TIM3, expressed in neutrophils. TIM3 expression levels increase after exposure to IL-8 or TNF-α and its binding with Gal9 would induce increased cytosolic calcium, causing neutrophil degranulation and thus increasing their cytotoxicity. In CF, TIM-3/Gal-9 signaling by neutrophils is disrupted in the airways due to proteolytic degradation of the receptor, as described in Bal of CF patients (SR76). As regards innate immunity in CF, PA elastase showed to induce the cleavage of monoclonal IgG and polyclonal IgG from CF patients, producing functional immune fragments effectively inhibiting phagocytosis of PA by the phagocytic cells present in this acute inflammatory lung injury (SR75). PA induces neutrophil extracellular trap (NET) formation in healthy donors and patients with CF. During inflammatory responses, activated human neutrophils and ECs secrete the antimicrobial peptide LL-37, a member of the cathelicidin family that facilitates NET formation. NET formation can be induced by signals that activate NADPH oxidase–dependent and –independent pathways.

To induce NET formation and proinflammatory cytokine release, PA engages CLEC5A, which is critical for PA–induced proinflammatory cytokine release and GSDMD cleavage in macrophages (SR99). In patients with COPD, there is an increasing amount of laminin deposition in the respiratory tract. The abundance of receptors in PA for laminin could be an important clinical element: in fact, PA isolates from the lung of patients with long term colonization bound significantly more laminin than isolates from patients with bacteremia (SR6). Gene expression analyses revealed changes in the abundance of mononuclear phagocytes (monocytes, macrophages, and dendritic cells) associated with CF severity, PA infection status, and disease progression (SR106). Multi-omics strategy enables to explore important signaling pathways to better understand the regulatory mechanisms following host infection with PA.

In mouse lung tissues, an alteration of chromatin accessibility contributes to the differential expression of genes associated with PA-infected lung tissues that have an important function in immunological regulation and in the prevention of microbial infection, such as Cxcl3, Il1r2, and Acod1. In the Reactome database, it was found that in PA-infected alveolar macrophages, key upregulated differentially expressed genes (DEGs) associated with differentially accessible regions (DARs) are linked to Cytokine signaling and Interleukin signaling in the immune system. In addition, they are also responsible for the regulation of the actin cytoskeleton, the JAK/STAT signaling pathway and the MAPK signaling pathway. Proteomics reveal key proteins in PA-infected lung tissues that act as biomarkers and determinants of inflammation and disease progression. Omics changes in Stat1 and Stat3 in mouse lung tissues and alveolar macrophages suggest these transcription factors play vital roles in immune defense and balance during PA infection (SR90).

Among these, some not only act as biomarkers of infection and inflammation but are major determinants of inflammatory status and disease progression: the expression of Ccl3 (chemokine C-C motif ligand 3) distinguishes *S. pneumoniae* infection from that from PA (SR89). Finally, omics changes in the transcription factors Stat1 and Stat3 in mice lung tissues and alveolar macrophages during PA infection, suggest that Stat1 and Stat3 may play vital roles in host immune defense and immune balance. It is found that PA genes encoded proteins associated with outer-membrane vesicles (OMV) are significantly up-regulated during infection, indicating the importance of OMV-associated proteins during acute respiratory infection; the same goes for genes encoded for essential amino acids, for genes involved in stress response, genes encoding the T3SS secreted effectors, virulence factors, and genes involved in flagellar biosynthesis. Regarding the host, the expression of genes encoding cytokines and chemokines are significantly increased during infection, as well as the genes encoding FAS and TNFR2 together with genes whose products are involved in TNF and NFκB signaling (SR93). Investigating the transcriptome profile of PA in a murine acute pneumonia model, a regulatory gene pvrA (PA2957, *Pseudomonas* virulence regulator A) was found to contribute to bacterial virulence and is upregulated during infection. Differential gene expression analysis shows that these changes in PA and in host gene expression occur not only during respiratory infection, but also during blood infection (SR94). Single-cell RNA sequencing (scRNA-seq) enables identification of the immune cell atlas in PA-infected lungs under acute and chronic conditions. PA infection alters the composition, distribution, and expression patterns of immune cell compartments critical for host defense. In acute infection, CD4+ naïve T cells, CD8+ naïve T cells, Cxcl2+ B cells, inflammatory monocytes, and neutrophils expand significantly, consistent with human studies. Neutrophils also increase in chronic infections. Additionally, interstitial macrophages (IMs) increase notably, while alveolar macrophages (AMs) decrease significantly in chronic infection. Two subtypes of AMs exhibit distinct roles in combating chronic PA infection, with IMs showing high expression of genes related to macrophage activation and antibacterial activity. Macrophages and neutrophils engage in dynamic cellular crosstalk during PA infection (SR105). The metabolomic analysis allowed to find that PA infection resulted in inhibition of the super pathway of quinolone, alkylquinolone biosynthesis, and 2-heptyl-3- hydroxy-4(1H)-quinolone biosynthesis of gut microorganisms in mice, proving that PA infection resulted in weakened anti-infection or anti-inflammatory ability of gut symbiotic bacteria in mice. Further, microbial functional analysis reveals that these changes in biosynthesis pathways are significantly decreased in the gut microbiota of mice model groups, suggesting that the gut microbiota in mice with pneumonia is disturbed, accompanied by a decrease in microbial diversity, number, and function changes (SR98).

**2. Bloodstream and systemic infection**

The vascular endothelium is a primary target for numerous human pathogenic bacteria and their virulence factors. When bacteria enter the bloodstream, they can evade immune defenses and spread throughout the body. This process can lead to alterations or disruptions of the endothelial barrier, triggering severe pathological events within the vasculature (SR57). In septic shock, TREM-1 (Triggering Receptor Expressed on Myeloid cells-1), an immunoreceptor expressed on neutrophils, monocytes/macrophages, and endothelial cells, amplifies the inflammatory response. TREM-1 deletion protected mice during septic shock by modulating inflammatory responses. The design of endothelium-specific TREM-1 inhibitors may prove interesting in preventing endothelial dysfunction (SR4). PA-LOX shows a specific cytotoxic activity towards human erythrocytes, permeabilizing red blood cell membranes. This could be related to the oxidation of membrane lipids. Since the peculiar capacity of PA-LOX to oxidize plasma membrane lipids of both human erythrocytes and human alveolar epithelial cells, suggests the enzyme might function as virulence factor during PA infections (SR45). Analysis of plasma from recent-onset Type 1 diabetes (RO T1D) patients and CF patients chronically infected with PA reveals that CF sera induce the transcription of IRF1, GIMAP1, GIMAP5, TLR10, IL32, CCL5, CD40, IKZF1, IL15, and IL16 in PBMCs. Conversely, CF sera downregulate many biological processes and IL-1 regulated genes involved in immune recognition and response, including PTGS2, CCL2, IRAK3, IL1B, and IL1R1, which are upregulated in T1D sera. This suggests distinct immune modulation patterns between CF and T1D sera (SR88). In order to identify the ability of PA to adapt and survive within the blood of severely burned patients during systemic infection, RNAseq analysis shows that the expression of QS genes is repressed, whereas the expression of pqsH, which codes for the FAD-dependent mono-oxygenase, increases. These results suggest that the QS systems may not be essential for the survival of PA in the blood during systemic infection (SR92). In a murine thermal injury model, metabolomic analysis identified specific biomarkers for sepsis caused by PA dissemination from infected burn wounds. PA infection alters the blood metabolome by consuming certain metabolites, decreasing their levels, and increasing the concentration of others. Using GC-TOF-MS and enrichment analysis, increased levels of pyrimidine metabolism-related metabolites (thymidine, thymine, uridine, and uracil) were found in the blood of septic thermally-injured mice (SR96). Further, a murine model of polytrauma and hemorrhagic shock that emulates severe, major human trauma, determined that the miR expression pattern in bone marrow HSPCs after polytrauma and pneumonia is markedly dissimilar in old versus young/adult mice (SR97). Transcriptomic analysis of the BSI-associated PA clinical isolates showed a high-level expression of cell-surface signaling (CSS) system Hxu, while a comparative genomic analysis of these isolates shows that a mutation in the rnfE gene is responsible for the elevated expression of the Hxu-CSS pathway (SR100). Deletion of the hxuIRA genes in PA reduces bloodstream infection (BSI) capability, whereas overexpression of hxuIRA genes enhances BSI in a murine sepsis model. Additionally, hemoglobin, haptoglobin, hemopexin, and transferrin in blood plasma can activate the Hxu system. This suggests that the Hxu system serves as a significant signal transduction pathway contributing to the adaptive pathogenesis of PA in BSI (SR100). From the results deriving from gene expression and gene regulatory processes, it emerged that among-individual differences can contribute to disease risk after pathogen exposures. Single-cell profiling of immune cells upon PA stimulation shows that myeloid cells (monocytes and DCs) exhibit the highest number of differential expression genes, whereas both CD4+ and CD8+ T cells show the fewest DE genes. This is consistent with the fact that innate immune cells are the first responders during pathogen stimulation. Further, some pathways, such as IL-1 signaling, were clearly enriched at a specific time-point of the infection (SR104).

**ABBREVIATIONS**

PA: *Pseudomonas aeruginosa*

ECs: Epithelial Cells

CTFR: cystic fibrosis transmembrane conductance regulator

CF: cystic fibrosis

c-JNK: c-Jun NH2-terminal kinase

MAPK: mitogen-activated protein kinase

IPF: idiopathic pulmonary fibrosis

PLA2: phospholipase A2

MMP7: matrix metalloproteinase 7

HSPGs: chains of heparan sulfate proteoglycans

AP: apical surface of polarized epithelium

BL: basolateral surface of polarized epithelium

QS: Quorum Sensing Genes

C4-HSL: N-butyryl-L-homoserine lactone

T3SS: Type III secretion system

GAP: GTPase-activating protein

AJ: apical junctions

TJ: tight junctions

ECM: extracellular matrix

OMV: outer-membrane vescicles

NET: neutrophil extracellular trap

CLEC5A: myeloid C-type lectin domain family 5-member A

GSDMD: gasdermin D

COPD: Chronic obstructive pulmonary disease

BSI: Bloodstream infections

VAP: Ventilator-associated pneumonia

PC: phosphatidylcholine

IL-1: interleukin-1

DC: dendritic cells

CASP3: Caspase-3

3O-C12-HSL: N-(3-oxododecanoyl)-l-homoserine lactone

A4GALT: Lactosylceramide 4-alpha-galactosyltransferase

ACTB: Actin, cytoplasmic 1

ADAM10: Disintegrin and metalloproteinase domain-containing protein 10

ALDOA: Fructose-bisphosphate aldolase A

algX: Alginate biosynthesis protein AlgX

ambB: AMB antimetabolite synthase AmbB

ambE: AMB antimetabolite synthase AmbE

APOE: Apolipoprotein E

aprA: Serralysin

AsialoGM1: Gangliotetraosylceramide (Galβ1,2GalNAcβ1,4Galβ1,4Glcβ1,1Cer)

azu: Azurin

C1q: Complement C1q

C1q: Complement C1q

C2: Complement C2

C3: Complement C3

C4: Complement C4

C5: Complement C5

CAMP: Cathelicidin antimicrobial peptide mature LL-37

CASP8: Caspase-8

CBLB: E3 ubiquitin-protein ligase CBL-B

CCN1: CCN family member 1

CCN2: CCN family member 2

CD14: Monocyte differentiation antigen CD14

CDC25C: M-phase inducer phosphatase 3

CDC42: Cell division control protein 42 homolog

CDH1: Cadherin-1

CDH5: Cadherin-5

CFH: Complement factor H

CFTR: Cystic fibrosis transmembrane conductance regulator

COL1A1: Collagen

CRK-I: Adapter molecule crk ISOFORM 1

CRK-II: Adapter molecule crk ISOFORM 2

CRKL:  CRK-like (CRKL)

CTNNB1: Catenin beta-1

DEF1: Neutrophil defensin 1

dinB: DNA polymerase IV

DNAJC5: DnaJ homolog subfamily C member 5

eco: Ecotin

EF2: Elongation factor 2

ELANE: Human Leukocyte elastase

estA: Esterase EstA

eta: exotoxin A

exoS: Secreted exoenzyme S

exoT: exoenzyme T

exoU: exoU

exoY: Adenylate cyclase exoY

EZR: Ezrin

F2: Prothrombin

FGA: Fibrinogen alpha chain

FHR-1: Complement factor H-related protein 1

fliC: A-type flagellin

FN1: Fibronectin

FOS: Protein c-Fos

H1B: Histone protein 1B

H2AX: Histone H2AX

HAVCR2: T cell immunoglobulin and mucin-domain containing-3 (TIM-3 / HAVCR2)

HCM: Lipid bilayers cell membrane

IFNG: Interferon gamma

IGHG1: immunoglobulin G

IL-2: Interleukin-2

IL-8: Interleukin-8

impA: Immunomodulating metalloprotease

ITGB1: Integrin beta-1

JUN: Protein c-Jun

LAMA1: Laminin subunit alpha-1

lasB: Elastase

lasI: Acyl-homoserine-lactone synthase

lecA: PA-I galactophilic lectin

LGALS9: Galectin-9

Lipid: A: Lipid A lipopolysaccaride

lox: Linoleate 9/13-lipoxygenase

lpd3: Dihydrolipoyl dehydrogenase

LPS: lipopolysaccaride

LRP1: Prolow-density lipoprotein receptor-related protein 1

LTF: Lactotransferrin

MAPK1: Mitogen-activated protein kinase 1

MAPK14: Mitogen-activated protein kinase 14

MAPK3: Mitogen-activated protein kinase 3

MAPKAPK2: MAP kinase-activated protein kinase 2

MASP: Mannan-binding lectin serine protease

MRAS: Ras-related protein M-Ras

MSN: moesin

MUC1: Mucin-1

MUC5AC: Mucin-5AC

MYD88: Myeloid differentiation primary response protein MyD88

NEU1: Sialidase-1

NOX1: NADPH oxidase 1

OCLN: Occludin

oprD: Porin D

oprF: Outer membrane porin F

oprG: Outer membrane protein OprG

oprH: PhoP/Q and low Mg2+ inducible outer membrane protein H1

oprQ: OprE3, belonging to outer membrane porin (Opr) (TC 1.B.25) family

PA3923: DUF1302 domain-containing protein

Paf: Probable adhesion protein

Phospholipid: Phospholipid cell membrane

PIK3C3: Phosphatidylinositol 3-kinase catalytic subunit type 3

pilA: Type IV major pilin protein PilA

pilY1: Type IV pilus biogenesis factor PilY1

PIP4P2: Type 2 phosphatidylinositol 4,5-bisphosphate 4-phosphatase

PLAUR: Urokinase plasminogen activator surface receptor

PLG: Plasmin

PLG: Plasminogen

popB: Translocator outer membrane protein PopB

popD: Translocator outer membrane protein PopD

POT1: Protection of telomeres protein 1

prpL: Lysyl endopeptidase

Psl: exopolysaccharide (Psl) syntetetized by polysaccaride syntetase locus

pumA: Monooxygenase PumA

RABEP1: Rab GTPase-binding effector protein 1

RAC1: Ras-related C3 botulinum toxin substrate 1

RALBP1: RalA-binding protein 1

RASA1: Ras GTPase-activating protein 1

RDX: radixin

RHOA: Transforming protein RhoA

RIPK3: Receptor-interacting serine/threonine-protein kinase 3

RND1: Rho-related GTP-binding protein Rho6

RND2: Rho-related GTP-binding protein RhoN

RND3: Rho-related GTP-binding protein RhoE

SELPLG: P-selectin glycoprotein ligand 1

SERPINA1: Alpha-1-antitrypsin

SFN: 14-3-3 protein sigma

SFTPA1: Pulmonary surfactant-associated protein A1

SLPI: secretory leukoprotease inhibitor

SMPD1: Sphingomyelin phosphodiesterase

SOD1: Superoxide dismutase [Cu-Zn]

SPN: CD44 antigen (Sialophorin)

TF: Serotransferrin

TGFA: Protransforming growth factor alpha

TIRAP: Toll/interleukin-1 receptor domain-containing adapter protein

TJP1: Tight junction protein ZO-1

TLR2: Toll-like receptor 2

TLR4: Toll-like receptor 4

TLR5: Toll-like receptor 5

TNFRSF1A: Tumor necrosis factor receptor superfamily member 1A

TP53: Cellular tumor antigen p53

TREM1: Triggering Receptor Expressed On Myeloid Cells 1

TRPV4: Transient receptor potential cation channel subfamily V member 4

tuf: Elongation factor Tu (EF-Tu)

UBB: Polyubiquitin-B

VIM: vimentin

VTN: Vitronectin

vWF: Subendothelial matrix-specific protein  (VON WILDENBRAND FACTOR)

YWHAB: 14-3-3 protein beta/alpha

YWHAE: 14-3-3 protein epsilon

YWHAG: 14-3-3 protein gamma

YWHAH: 14-3-3 protein eta

YWHAQ: 14-3-3 protein theta

YWHAZ: 14-3-3 protein zeta/delta
